# Tumor suppressor genotype influences the extent and mode of immunosurveillance in lung cancer

**DOI:** 10.1101/2025.01.15.633175

**Authors:** Keren M. Adler, Haiqing Xu, Amy C. Gladstein, Valerie M. Irizarry-Negron, Maggie R. Robertson, Katherine R. Doerig, Dmitri A. Petrov, Monte M. Winslow, David M. Feldser

**Author notes:** Equal Contribution (Keren M. Adler, Haiqing Xu).

## Abstract

The impact of cancer driving mutations in regulating immunosurveillance throughout tumor development remains poorly understood. To better understand the contribution of tumor genotype to immunosurveillance, we generated and validated lentiviral vectors that create an epi-allelic series of increasingly immunogenic neoantigens. This vector system is compatible with autochthonous Cre-regulated cancer models, CRISPR/Cas9-mediated somatic genome editing, and tumor barcoding. Here, we show that in the context of KRAS-driven lung cancer and strong neoantigen expression, tumor suppressor genotype dictates the degree of immune cell recruitment, positive selection of tumors with neoantigen silencing, and tumor outgrowth. By quantifying the impact of 11 commonly inactivated tumor suppressor genes on tumor growth across neoantigenic contexts, we show that the growth promoting effects of tumor suppressor gene inactivation correlate with increasing sensitivity to immunosurveillance. Importantly, specific genotypes dramatically increase or decrease sensitivity to immunosurveillance independently of their growth promoting effects. We propose a model of immunoediting in which tumor suppressor gene inactivation works in tandem with neoantigen expression to shape tumor immunosurveillance and immunoediting such that the same neoantigens uniquely modulate tumor immunoediting depending on the genetic context.

**One Sentence Summary:** Here we uncover an under-appreciated role for tumor suppressor gene inactivation in shaping immunoediting upon neoantigen expression.

## INTRODUCTION

Our current understanding of cancer immune surveillance is colored by the immunoediting hypothesis, which proposes that the adaptive immune system imposes a selective pressure that drives a tumor’s evolution towards a more immune evasive state in response to neoantigens presented on cancer cells (*1, 2*). In the decades since the inception of the immunoediting hypothesis, multiple mechanisms have emerged to explain how cancer cells actively promote an immune suppressive tumor microenvironment that limits the immune system’s ability to identify and eliminate antigen-expressing cancer cells (*3–9*). Interestingly, the precise genetic determinants of malignant transformation itself (*i.e.* oncogene acquisition and tumor suppressor gene inactivation) can regulate cell-intrinsic immune modulatory pathways and even predict the response to immune checkpoint therapy in certain contexts (*10–19*). However, the extent to which cancer genotype shapes immune surveillance is only beginning to be understood, and the extent to which different genotypes increase or decrease immunoediting remains unclear.

Autochthonous cancer models that recapitulate tumor initiation and early expansion due to relevant oncogene activation and concomitant tumor suppressor gene inactivation represent tractable model systems to understanding gene-microenvironment interactions in a physiologically relevant context (*20, 21*). Recent advancements in CRISPR-mediated gene inactivation in somatic cells coupled with tumor barcoding have enabled the quantitative analysis of gene-microenvironment interactions on an unprecedented scale and with profound resolution to detect changes in tumor cell fitness (*22*). Here, we combine genetically engineered conditional mouse models with lentiviral-mediated somatic gene inactivation, neoantigen expression and tumor barcoding to better understand how inactivation of commonly mutated tumor suppressor genes modulates immunoediting. By expressing fixed neoantigens in different genetic contexts, we shift the focus away from the role of the neoantigen in immunoediting towards the importance of the genetic context in which the neoantigen arises. Our methodology is a resource for assessment of tumor genotype-immune phenotype interactions in a highly quantitative and relatively high throughput *in vivo* manner. Our study uncovers tumor genotype-dependencies that impact mechanisms of immune evasion as well as sensitivity or resistance to immune surveillance, thus highlighting a previously under-appreciated role for cancer driver mutations in shaping immunoediting.

## RESULTS

### A system to generate an epi-allelic series of immunogenicity in lung tumors *in vivo*

To model immunoediting during lung tumor formation, we used the *Kras^LSL-G12D/+^*(*K*) genetically engineered mouse where tumors are initiated via transduction of lung epithelial cells with lentiviral vectors expressing Cre recombinase (*20*). To create varying levels of immunogenicity, we generated a series of lentiviral vectors that express increasingly potent neoantigens. Poorly immunogenic tumors are initiated with Lenti:Cre which expresses only Cre recombinase. Mildly immunogenic tumors are initiated with Lenti:mCh/Cre which expresses mCherry and Cre. Finally, highly immunogenic tumors are initiated with Lenti:mCh-SIIN/Cre which expresses the MHC-I restricted peptide SIINFEKL linked to mCherry and Cre (Fig. 1A). Tumors were initiated in *K* mice with either Lenti:Cre, Lenti:mCh/Cre, or Lenti:mCh-SIIN/Cre, and analyzed 8, 12, and 16 weeks after tumor initiation (Fig. 1B). Mice transduced with each of the three vectors had numerous small adenomas 8 weeks after tumor initiation (Fig. 1C). As expected, tumors initiated with Lenti:Cre continuously expanded in size over time (Fig. 1, C and D). In contrast, tumors initiated with either Lenti:mCh/Cre or Lenti:mCh-SIIN/Cre expanded negligibly from 8 to 16 weeks. Immunohistochemical (IHC) staining for mCherry showed that the vast majority of tumors initiated with Lenti:mCh/Cre were mCherry^Positive^ throughout the 16 week time course (Fig. 1, E and F). In contrast, while the majority of tumors initiated with Lenti:mCh-SIIN/Cre (64%) were mCherry^Positive^ 8 weeks after tumor initiation, progressively fewer tumors expressed mCherry at 12 (39%) and 16 (2%) weeks after tumor initiation (Fig. 1,E and F).

**Fig. 1.**
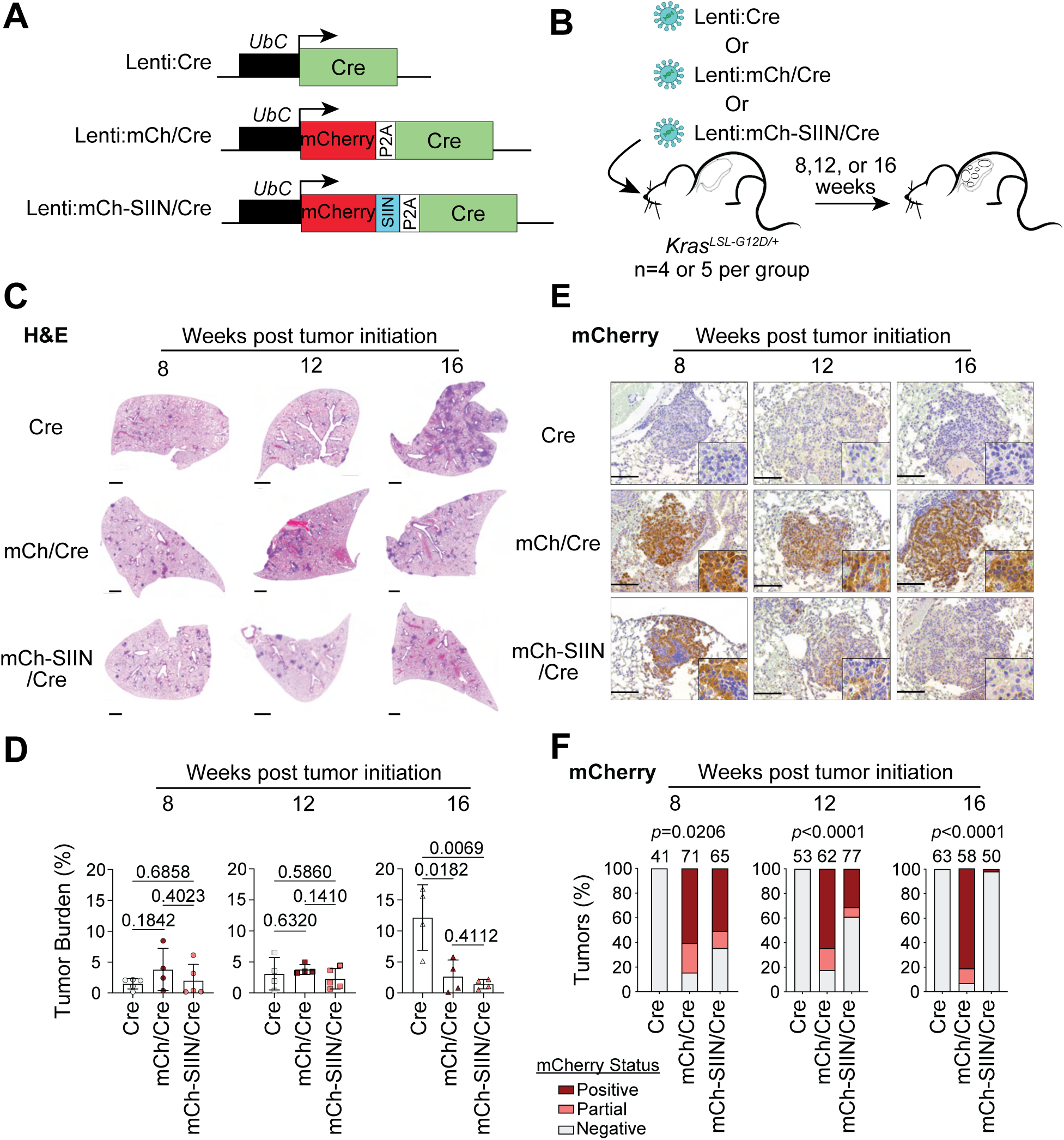
An epi-allelic series of lentiviral vectors to study tumor immunosurveillance in KRAS-driven lung tumors. **A.** Depiction of epi-allelic series of lenti-vectors used to initiate tumors. Top, Lenti:Cre: baseline vector expresses Cre recombinase from the UbC promoter. Middle, Lenti:mCh/Cre: moderately immunogenic vector expresses mCherry and Cre recombinase from the UbC promoter. Bottom, Lenti:mCh-SIIN/Cre: highly immunogenic vector expresses an mCherry-SIINFEKL fusion protein as well as Cre recombinase from the UbC promoter. **B.** Experimental outline. Tumors were initiated in *Kras^LSL-G12D/+^*mice using 100,000 pfu of Lenti:Cre, Lenti:mCh/Cre, or Lenti:mCh-SIIN/Cre. Lungs were harvested for sectioning at 8, 12, or 16 weeks post-transduction. **C.** Representative scans of H&E stained lungs from each experimental group. Tile scans were taken at 5x magnification. Scale bar is 1mm. **D.** Quantification of percent tumor burden for each vector at 8, 12, and 16 weeks. Statistical significance was determined using unpaired Student’s *t*-tests. Error bars represent mean ± standard deviation. For Lenti:Cre, n=5 mice at 8 weeks post tumor initiation and n=4 mice at 12 and 16 weeks post tumor initiation. For Lenti:mCh/Cre, n=4 mice at 8, 12, and 16 weeks post tumor initiation. For Lenti:mCh-SIIN/Cre, n=5 mice at 8 and 12 weeks post tumor initiation and n=4 mice at 16 weeks. **E.** Representative IHC images for mCherry at 8, 12, and 16 weeks after tumor initiation with Lenti:Cre, Lenti:mCh-Cre, or Lenti:mCh-SIIN/Cre. Insets are 3x magnified. Scale bar is 119um. **F.** Qualitative assessment of mCherry expression based on IHC from Figure 1E. Tumors were split into three groups based on mCherry expression status: positive, partial, and negative. Values were counted and graphed as percentages. Number of individual tumors analyzed for each group is indicated on graph. For Lenti:Cre, n=5 mice at 8 weeks, n=4 mice at 12 and 16 weeks. For Lenti:mCh/Cre, n=4 mice at 8, 12, and 16 weeks. For Lenti:mCh-SIIN/Cre, n=5 mice at 8 and 12 weeks, and n=4 mice at 16 weeks.

To determine the extent to which each immunogenic condition promotes immune cell infiltration, we performed IHC for CD45 and CD3 (Fig. 2, A and B). Tumors initiated with Lenti:Cre had relatively few infiltrating immune cells which remained constant throughout the 8, 12, and 16 week time points after tumor initiation (Fig. 2C). Tumors initiated with Lenti:mCh/Cre also had low level immune infiltration at 8 and 12 weeks after tumor initiation similar to that of Lenti:Cre. However, 16 weeks after tumor initiation, Lenti:mCh/Cre tumors had a dramatic and significant increase in the number of CD45^Positive^ and CD3^Positive^ immune cells (Fig. 2, C and D). In contrast, tumors initiated by Lenti:mCh-SIIN/Cre had high immune infiltration early that progressively increased over time such that by the 16 week time point CD3^Positive^ cells were the dominant cell type in some tumors (Fig. 2, C and D). The persistence of immune infiltration in tumors initiated with Lenti:mCh/Cre or Lenti:mCh-SIIN/Cre at the 16 week time point is consistent with their constricted expansion compared to tumors initiated with Lenti:Cre (Fig. 1, C and D). Taken together, these data suggest that while mCherry expression alone elicits an immune response strong enough to restrict tumor outgrowth, it is insufficiently immunogenic to select for the outgrowth of cells with neoantigen silencing. However, mCherry-SIIN expression is sufficiently potent to drive rapid selection of tumor cells that have silenced neoantigen expression. Thus, Lenti:Cre, Lenti:mCh/Cre, and Lenti:mCh-SIIN/Cre represent an epi-allelic series of neoantigens that can be expressed within autochthonous lung tumors to model varying degrees of immunogenicity *in vivo*.

**Fig. 2.**
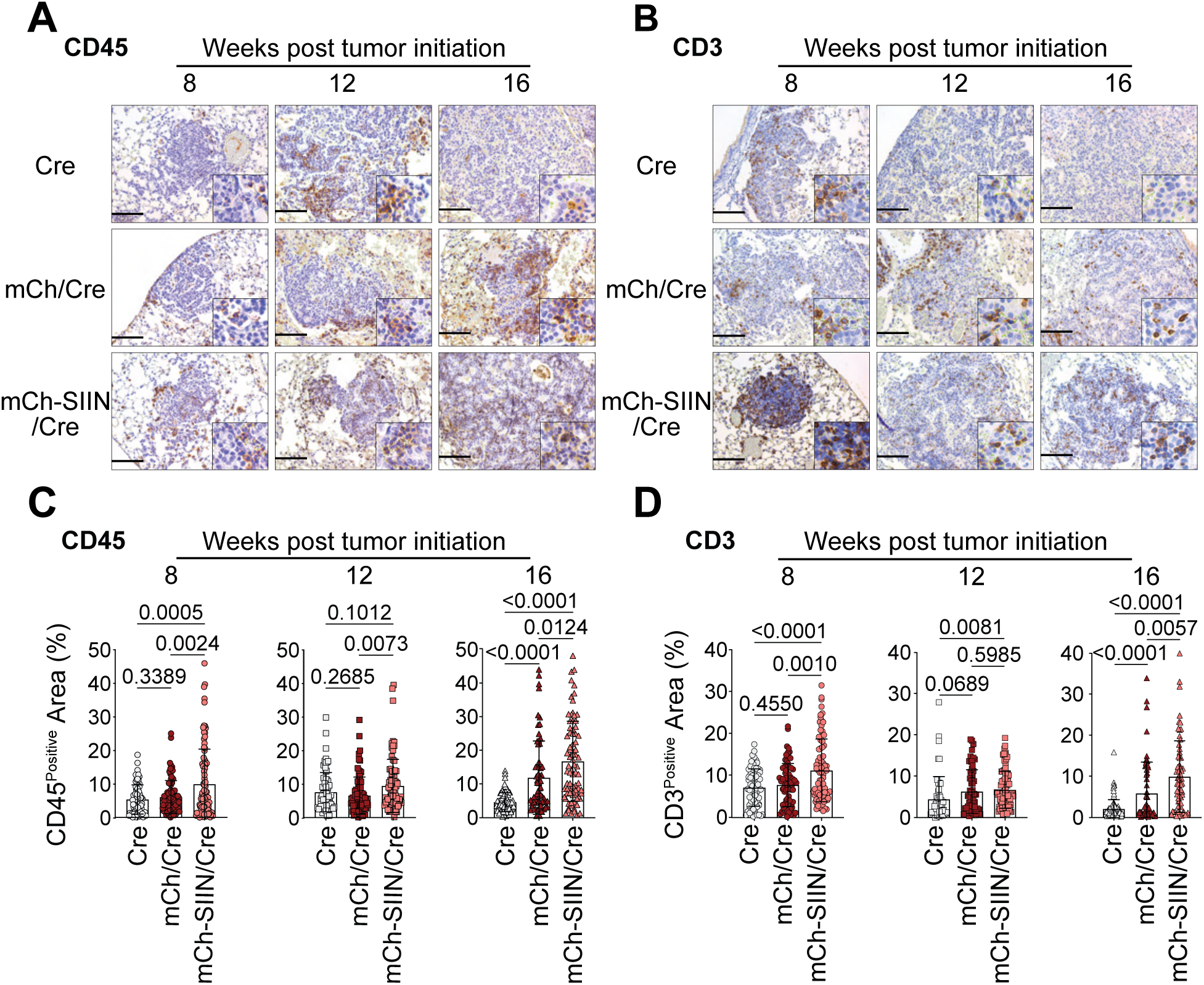
SIINFEKL expression induces an immune response in KRAS-driven lung tumors. **A, B.** Representative 20x IHC for the immune markers CD45 (A) and CD3 (B) at 8, 12, or 16 weeks post-tumor initiation in tumors initiated by Lenti:Cre, Lenti:mCh/Cre, or Lenti:mCh-SIIN/Cre. Insets are 3x magnified. Scale bar is 119um. **C.** Quantification of CD45 IHC shown in Figure 2A using percent positive area. Statistical significance was determined using unpaired Student’s *t*-tests. Error bars represent mean ± standard deviation. For Lenti:Cre, n=77 tumors from n=5 mice at 8 weeks, n=71 tumors from n=4 mice at 12 weeks, and n=80 tumors from n=4 mice at 16 weeks. For Lenti:mCh/Cre, n=84 tumors from n=4 mice at 8 weeks, n=81 tumors from n=4 mice at 12 weeks, and n=65 tumors from n=4 mice at 16 weeks. For Lenti:mCh-SIIN/Cre, n=93 tumors from n=5 mice at 8 weeks, n=84 tumors from n=5 mice at 12 weeks, and n=80 tumors from n=4 mice at 16 weeks. **B.** Quantification of CD3 IHC shown in Figure 2B using percent positive area. Statistical significance was determined using unpaired Student’s *t*-tests. Error bars represent mean ± standard deviation. For Lenti:Cre, n=71 tumors from n=5 mice at 8 weeks, n=52 tumors from n=4 mice at 12 weeks, and n=86 tumors from n=4 mice at 16 weeks. For Lenti:mCh/Cre, n=72 tumors from n=4 mice at 8 weeks, n=62 tumors from n=4 mice at 12 weeks, and n=61 tumors from n=4 mice at 16 weeks. For Lenti:mCh-SIIN/Cre, n=86 tumors from n=5 mice at 8 weeks, n=67 tumors from n=5 mice at 12 weeks, and n=66 tumors from n=4 mice at 16 weeks.

### Tumor suppressor genotype differentially regulates immunoediting

To alter tumor suppressor genotype in the context of neoantigen expression, we incorporated U6 promoter-driven single guide RNAs (sgRNAs) into the Lenti:mCh/Cre and Lenti:mCh-SIIN/Cre vectors (Fig. 3A). We initiated tumors in *Kras^LSL-G12D/+^;Rosa26^LSL-Cas9::GFP/LSL-Cas9::GFP^* (*K;Cas9*) mice with Lenti:mCh/Cre and Lenti:mCh-SIIN/Cre vectors that express sgRNAs targeting one of three commonly inactivated tumor suppressor genes in lung adenocarcinoma (*Lkb1*, *Setd2*, or *Rb1*) or express an inert non-targeting sgRNA to generate control tumors driven solely by oncogenic KRAS (TS^WT^) (Fig. 3B). To best capture changes in tumor outgrowth associated with potent neoantigen expression, lungs were analyzed 12 weeks after tumor initiation. As expected, *Lkb1^KO^*, *Setd2^KO^*, or *Rb1^KO^* significantly increased tumor growth compared to TS^WT^ tumors initiated by Lenti:mCh/Cre (Fig. 3, C and D). Unexpectedly, *Lkb1^KO^*, *Setd2^KO^*, or *Rb1^KO^* did not significantly increase tumor burden compared to TS^WT^ tumors when initiated with Lenti:mCh-SIIN/Cre. Thus, strong neoantigen expression effectively limits the growth promoting effects of inactivating these tumor suppressor genes (Fig. 3, C and D).

**Fig. 3.**
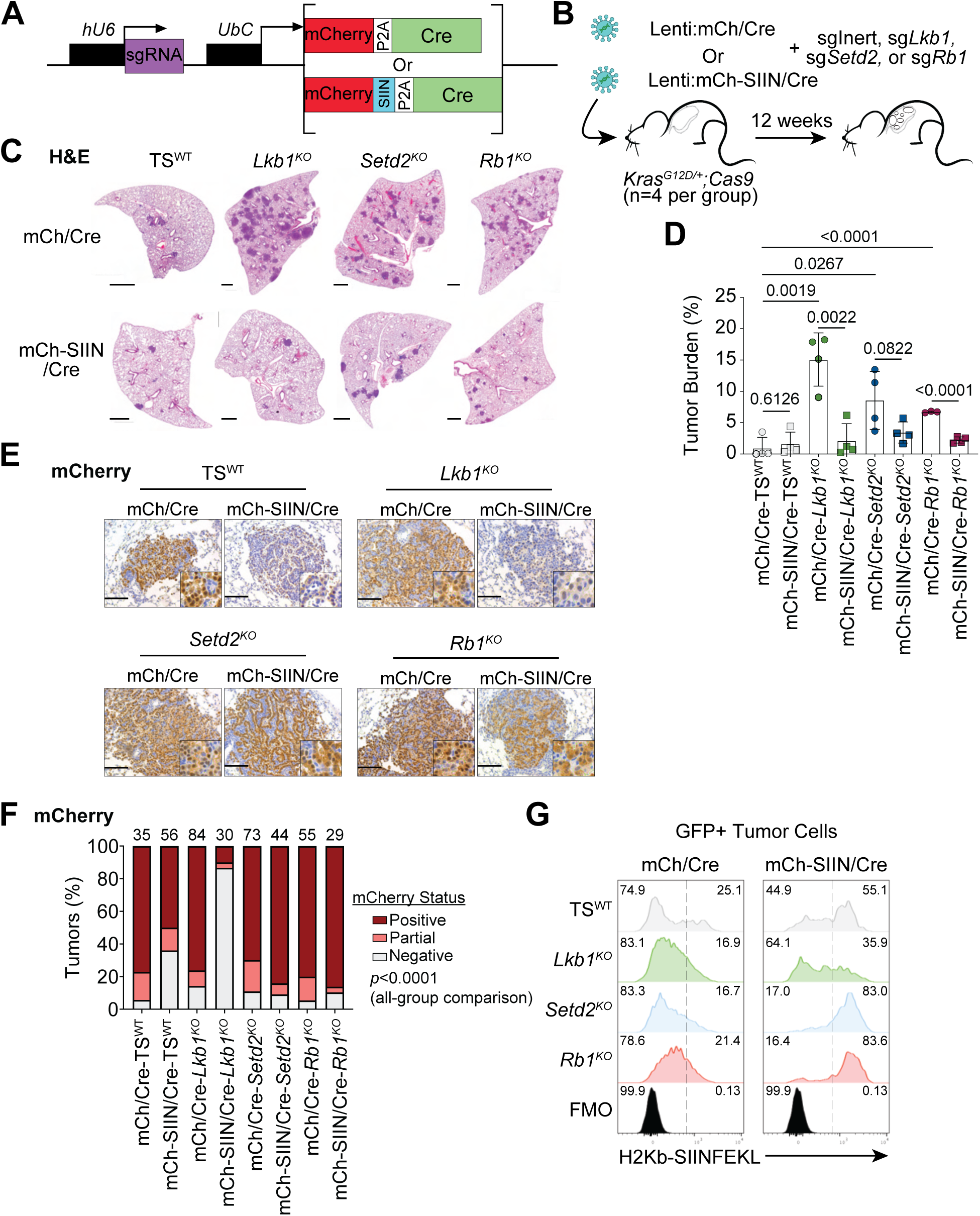
Tumors with *Setd2^KO^*or *Rb1^KO^*, but not *Lkb1^KO^*, maintain SIINFEKL expression and presentation 12 weeks after tumor initiation. **A.** Design of epi-allelic series of lenti-vectors used to simultaneously initiate tumors and knockout a gene of interest. Design is similar as to what is depicted in Figure 1A, with the addition of a human U6 promoter driving the expression of a sgRNA of interest. **B.** Experimental scheme. Tumors were initiated in *K;Cas9* mice with 60,000 pfu of Lenti:mCh/Cre or Lenti:mCh-SIIN/Cre containing sgRNAs targeting the tumor suppressors *Lkb1*, *Setd2*, or *Rb1*, in addition to an inert, non-targeting sgRNA. Mice were harvested 12 weeks after tumor initiation. **C.** Representative H&E stained lung lobes from each experimental group. Scans were taken at 5x magnification. Scale bar is 1mm. **D.** Percent tumor burden quantification for all experimental groups. Statistical significance was determined using unpaired Student’s *t*-tests. Error bars represent mean ± standard deviation. n=4 mice for all experimental groups except for mCherry-Rb1^KO^ (n=3). **E.** Representative IHC images of mCherry for each experimental group. Insets are 3x magnified and scale bar is 119um. **F.** Qualitative assessment of mCherry expression in each experimental group. Each tumor image was divided into one of three categories based on its mCherry expression level: positive, partial, or negative. Significance across all groups was determined using a chi-squared test. Number of individual tumors analyzed for each group is indicated on graph. n=4 mice for all groups except for, mCh/Cre-*Rb1^KO^*, where n=3 mice. **G.** Histogram plots showing H2Kb-SIINFEKL presentation measured by flow cytometry across each genotype specified. Plots are separated based on initiating vector and are normalized to mode. Dotted line separates positive and negative populations and value in the upper corner indicates the percentage of cells in that population.

In each genetic and immunogenic context, we assessed mCherry expression via IHC. Consistent with our initial IHC analysis (Fig. 1, E and F), the majority of TS^WT^ tumors initiated by Lenti:mCh/Cre were mCherry^Positive^, as were *Lkb1^KO^*, *Setd2^KO^*, or *Rb1^KO^* tumors initiated by Lenti:mCh/Cre (Fig. 3, E and F). Also consistent with our initial data (Fig. 1, E and F), a lower percentage of TS^WT^ tumors initiated by Lenti:mCh-SIIN/Cre were mCherry^Positive^, again suggesting a selection against mCherry-SIIN expression. However, the propensity to select for mCherry-SIIN silencing in cancer cells was distinct amongst *Lkb1^KO^*, *Setd2^KO^*, or *Rb1^KO^*tumors. *Lkb1^KO^* tumors were almost all mCherry^Negative^, while *Setd2^KO^* and *Rb1^KO^* tumors were similarly mCherry^Positive^ to those initiated by Lenti:mCh/Cre. Additionally, we directly measured mCherry expression and MHC-I/SIINFEKL antigen presentation on cancer cells via flow cytometry of tumor cells (Fig. S1A). In each group, MHC-I expression was uniformly high but SIINFEKL presentation and mCherry expression were consistent with expression observed via IHC (Fig. 3, E to G, Fig. S2). These data point to a selection of cancer cells with mCherry-SIIN silencing rather than a general loss of MHC-I expression or lack of antigen presentation. Taken together, these data suggest that while mCherry-SIIN expression promotes immunoediting across different tumor suppressor genotypes, the mechanism of immune evasion differs between *Lkb1^KO^*, *Setd2^KO^*, and *Rb1^KO^* cancer cells.

### Tumor suppressor genotype differentially regulates immune responses to neoantigens

To gain insight into whether tumor suppressor genotype impacts the immune response to defined neoantigens, we quantified immune infiltration using IHC in lung tumors initiated with each Lenti:mCh/Cre or Lenti:mCh-SIIN/Cre vector. Across every tumor suppressor genotype, there was a significant increase in CD45^Positive^, CD3^Positive^ and CD8^Positive^ immune cells in tumors initiated by Lenti:mCh-SIIN/Cre compared to tumors initiated by Lenti:mCh/Cre, indicating that mCherry-SIIN elicits a more potent anti-tumor immune response than mCherry alone regardless of tumor suppressor genotype (Fig. 4, A to C and Fig. S3, A to C). However, infiltration of immunosuppressive cell types in response to SIINFEKL expression varied dramatically based on tumor suppressor genotype. Compared to TS^WT^ tumors initiated by Lenti:mCh-SIIN/Cre, FoxP3^Positive^ regulatory T cell (T_reg_) infiltration was significantly higher in *Lkb1^KO^* and *Rb1^KO^* tumors but not in *Setd2^KO^* tumors (Fig. 4, A and C, Fig. S3D). Ly6G^Positive^ neutrophil infiltration was higher in the *Rb1^KO^*tumors but lower in *Setd2^KO^* tumors (Fig. 4, A and C, Fig. S3E). Arginase I^Positive^ cell infiltration, likely myeloid-derived suppressor cells (MDSCs), was higher in *Setd2^KO^* tumors but lower in *Lkb1^KO^* tumors (Fig. 4, A and C, Fig. S3F). Thus, this epi-allelic series of neoantigenic vectors facilitates the functional assessment of tumor genotypes within diverse immunogenic contexts and provides insight into mechanisms of genotype-dependent immune surveillance.

**Fig. 4.**
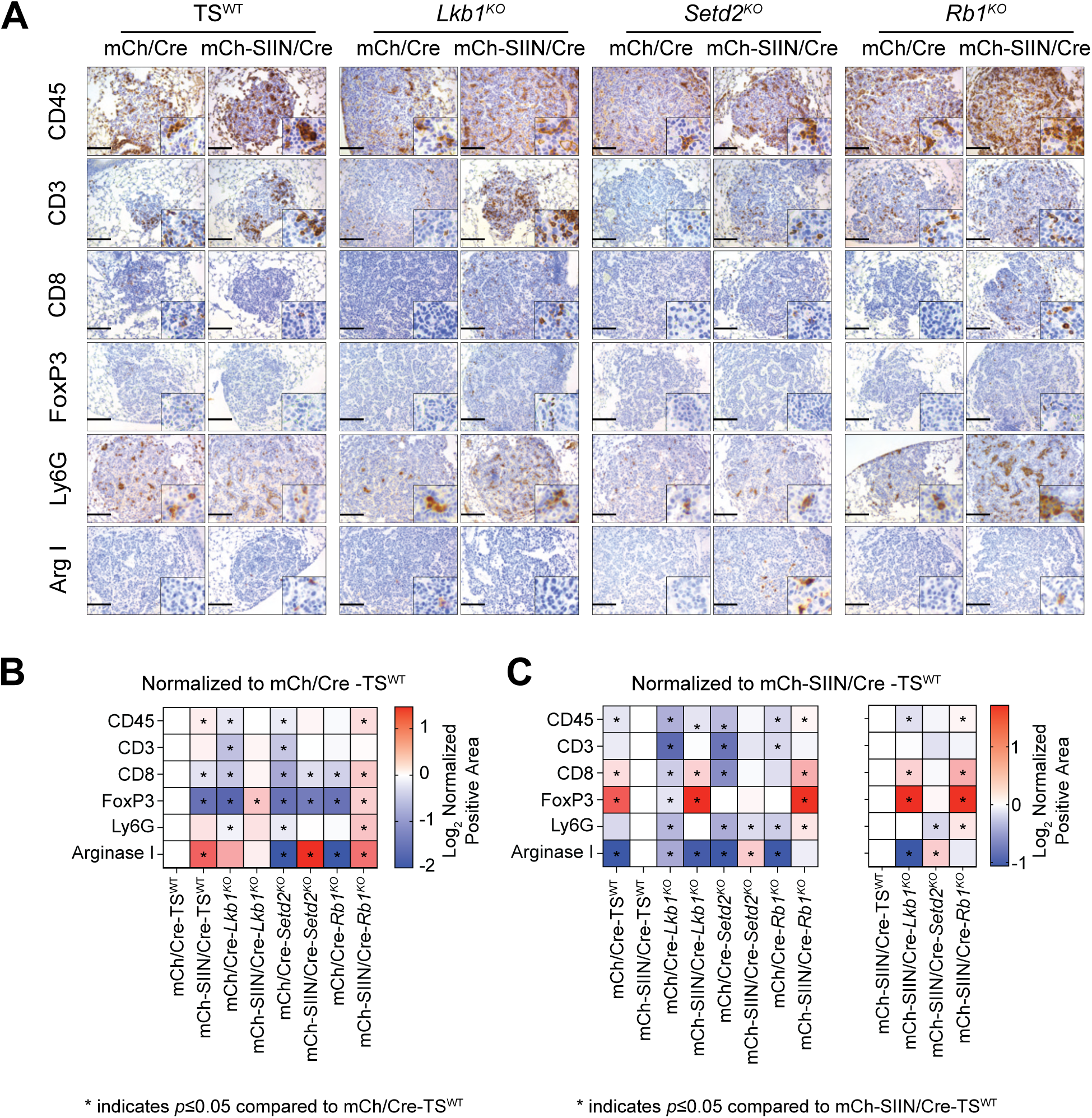
Tumor suppressor genotype differentially modulates immune responses in the context of potent antigen. **A.** Representative 20x IHC images for the immune markers CD45, CD3, CD8, FoxP3, Ly6G, and Arginase I for all 8 experimental groups outlined in Figure 3B. Insets are 3x magnified. Scale bar is 119um. **B.** Heat map depicting Log_2_ positive area for each IHC marker. Data are normalized to Lenti:mCh/Cre-TS^WT^. Asterisk indicates *p*-value<0.05 by Student’s *t-*test. **C.** Heat map depicting Log_2_ positive area for each IHC marker across all 8 experimental groups (left) or all 4 Lenti:mCh-SIIN/Cre groups (right). Data are normalized to Lenti:mCh-SIIN/Cre-TS^WT^. Asterisk indicates *p*-value<0.05 by Student’s *t-*test.

### Parallel analysis of tumor genotype across multiple immunogenic contexts via tumor barcoding

To assess the impact of multiple tumor suppressor genotypes in parallel across different immunogenic contexts in a highly quantitative manner, we generated sgRNA-expressing versions of Lenti:Cre, Lenti:mCh/Cre, and Lenti:mCh-SIIN/Cre for tumor barcoding coupled with high-throughput barcode sequencing (Tuba-seq, Fig. 5A). Tuba-seq is based on lentiviral-mediated integration of barcode sequences into the genome of the initiating cell of each clonal tumor in the lung. To enable pooling of different immunogenic vectors and the generation of different tumor suppressor genotypes, we inserted a 4 nucleotide vector ID (vID which is unique to each immunogenic vector) and a diverse tumor barcode (BC which is unique to each clonal tumor) within the 5’ region of the U6 promoter directly preceding the sgRNA (*22–26*). Thus, amplification and high-throughput sequencing of the vID-BC-sgRNA region from bulk tumor-bearing lung DNA can quantify the number of cancer cells in each tumor clone (BC reads), its genotype (sgRNA), and which immunogenic vector it contains (vID).

**Fig. 5.**
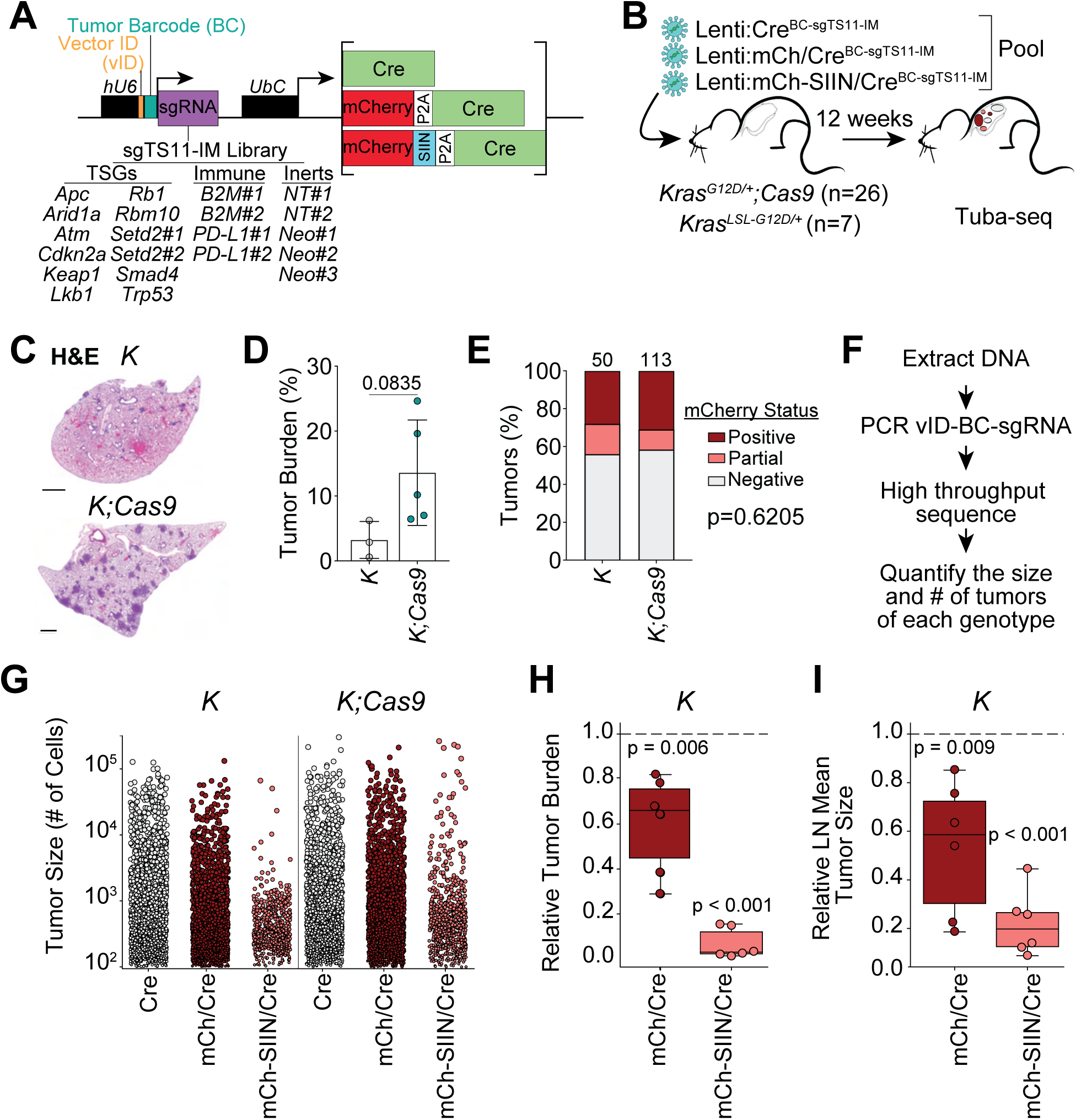
Multiplexed somatic genome editing and tumor barcoding uncover the impact of tumor suppressor gene inactivation on immunosurveillance. **A.** Design of epi-allelic series of lenti-vectors used for Tuba-seq screen. These lenti-vectors include a 4nt vector ID, a 10nt tumor barcode, as well as a library of sgRNAs targeting frequently mutated tumor suppressor genes in lung adenocarcinoma, immunomodulatory genes, as well as inert, non-targeting sgRNAs. **B.** Experimental design. Tumors were initiated in either *K* (n=7) or *K;Cas9* (n=26) mice using a pool of Lenti:Cre, Lenti:mCh/Cre, and Lenti:mCh-SIIN/Cre each containing the sgRNA library depicted in Figure 5A. Each *K* mouse received 100,000 pfu per virus (300,000 pfu total virus per mouse), and each *K;Cas9* mouse received 60,000 pfu per virus (180,000 pfu total virus per mouse). **C.** Scans of a representative H&E stained lung lobe from a *K* or *KC* mouse at 5x magnification. Scale bar is 1mm. **D.** Quantification of tumor burden for *K* (n=3) and *K;Cas9* (n=5) mice. Statistical significance was determined using an unpaired Student’s *t*-test. Error bars represent mean ± standard deviation. **E.** Qualitative assessment of mCherry expression in *K* (n=3) and *K;Cas9* (n=5) mice based on IHC from Supplementary Figure 5D. Number of individual tumors analyzed for each group is indicated on graph. Significance was determined using a chi-squared test. **F.** Flow chart summarizing the Tuba-seq pipeline from sacrificing mice to analyzing sequencing results. **G.** Jitter plot of all barcoded tumors across each immunogenic context (Cre^BC-sgTS11-IM^, mCh/Cre^BC-sgTS11-IM^, and mCh-SIIN/Cre^BC-sgTS11-IM^) in *K* mice and all barcoded TS^WT^ tumors in *K;Cas9* mice. Tumors were analyzed using the “Normal method” (Fig. S6A). **H and I.** Tumor burden (H) or LN mean tumor size (I) of mCh/Cre^BC-sgTS11-IM^ and mCh-SIIN/Cre^BC-sgTS11-IM^ tumors relative to Cre^BC-sgTS11-IM^ tumors across *K* mice. Tumors were analyzed using the “Normal method” (Fig. S6A). One-sample t-tests were used to determine if the ratios were significantly different from 1.

We assembled sgRNA libraries that target 11 tumor suppressor genes that are commonly inactivated in human lung adenocarcinoma (TS11, Fig. 5A). Additionally, we included sgRNAs targeting the immunomodulatory (IM) genes beta-2-microglobulin (*B2M*) and *CD274* (*PD-L1*) as controls that should have opposite effects on tumor outgrowth in a highly immunogenic context, as well as inert/non-targeting (sgInert) control sgRNAs (21 total sgRNAs in each vector backbone; Fig. 5A). A pool of all barcoded sgRNAs was cloned into each vector backbone to generate Lenti:Cre^BC-sgTS11-IM^, Lenti:mCh/Cre^BC-sgTS11-IM^, and Lenti:mCh-SIIN/Cre^BC-sgTS11-IM^ (Fig. 5A, Fig. S4A, and Methods). We transduced *K* and *K;Cas9* mice with a pool of the Lenti:Cre^BC-sgTS11-IM^, Lenti:mCh/Cre^BC-sgTS11-IM^, and Lenti:mCh-SIIN/Cre^BC-sgTS11-IM^ vectors (63 vectors total) and analyzed lungs 12 weeks after tumor initiation (Fig. 5, B and C, Fig. S4B). As expected, due to inactivation of potent tumor suppressors, tumor burden and individual tumor area in *K;Cas9* mice were larger than *K* mice (Fig. 5, C and D, Fig. S4, B and C). A subset of tumors were mCherry^Positive^ consistent with mCherry expression in a fraction of tumors initiated by Lenti:mCh/Cre^BC-sgTS11-IM^ and Lenti:mCh-SIIN/Cre^BC-sgTS11-IM^ (Fig. 5E, Fig. S4, B and D). The vID-BC-sgRNA region was PCR amplified from genomic DNA isolated from tumor bearing lungs, followed by high-throughput sequencing and Tuba-seq analysis (Fig. 5F, Fig. S5, A and B, and Methods).

To establish the baseline effects of expressing mild or strong neoantigens on immunosurveillance in TS^WT^ tumors, we quantified the relative impact of mCherry or mCherry-SIIN expression on tumor burden and tumor size compared to poorly immunogenic tumors initiated by Lenti:Cre^BC-sgTS11-IM^. Tumors initiated with either Lenti:mCh/Cre^BC-sgTS11-IM^ or Lenti:mCh-SIIN/Cre^BC-sgTS11-IM^ had a significantly lower tumor burden and smaller log normal mean tumor size (Fig. 5, G to I, Fig. S5, C to F). Moreover, Lenti:mCh-SIIN/Cre^BC-sgTS11-IM^ reduced tumor burden and tumor size more than Lenti:mCh/Cre^BC-sgTS11-IM^. Taken together, these findings provide additional quantitative support for the increasing tumor immunogenicity of each lentiviral vector and establishes Tuba-seq as a method to quantitatively and precisely assess the impact of neoantigen expression on immunosurveillance in different immunogenic contexts.

### CRISPR-mediated inactivation of mediators of immune surveillance predictably impacts tumor size

*B2M* is an essential component of the major histocompatibility complex type I (MHC-I). Loss of MHC-I though *B2M* inactivation drives resistance to immune checkpoint therapy and is predicted to also facilitate resistance to immunoediting in neoantigen expressing tumors (*7–9*). However, MHC-I expression is a determinant of ‘*self’* and inhibits innate immune responses that cull MHC-I negative cells (*27, 28*). As such, *B2M^KO^* may have complex effects on tumor number and tumor size *in vivo*. Indeed, relative to Inert (TS^WT^) tumors, *B2M^KO^* greatly reduced the size of poorly immunogenic Cre tumors (Fig. 6A, Fig. S6, S7, A and B). However, *B2M^KO^*slightly reduced the size of the mildly immunogenic mCherry tumors, and greatly increased the size in the highly immunogenic mCherry-SIIN tumors. Furthermore, *B2M^KO^* increased tumor initiation across all immunogenic conditions (Fig. S7C). These data suggest that while loss of self is generally detrimental to tumor growth, the selective advantage of losing SIINFEKL presentation outweighs the innate immune response induced by lack of MHC-I. In contrast, inactivation of *PD-L1* should sensitize tumors to immunoediting. *PD-L1^KO^*had no effect on the size of Cre tumors, and only a mildly negative effect on mCherry tumors. However, *PD-L1^KO^* greatly reduced the size and number of highly immunogenic mCherry-SIIN tumors indicative of enhanced immunoediting (Fig. 6A, Fig. S7, A to C). These results further underscore the graded immunogenicity of Lenti:Cre^BC-sgTS11-IM^, Lenti:mCh/Cre^BC-sgTS11-IM^, and Lenti:mCh-SIIN/Cre^BC-sgTS11-IM^ within autochthonous tumors and demonstrate our ability to discern complex effects of tumor genotype on the susceptibility to immune surveillance.

**Fig 6.**
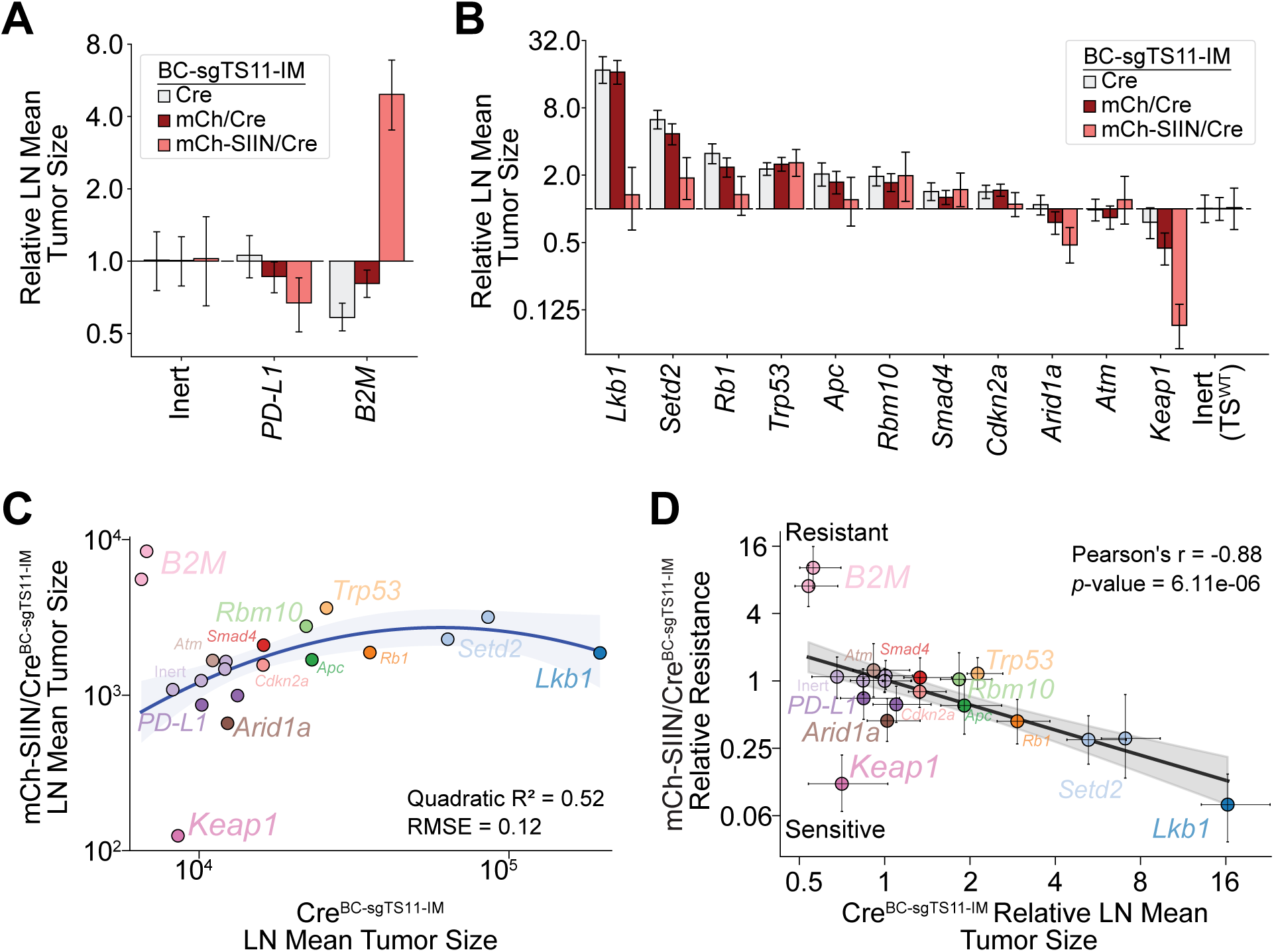
Highly proliferative tumor suppressor genotypes exhibit lower resistance to immunosurveillance. **A.** Quantification of relative LN mean tumor size for the TS^WT,^ *PD-L1^KO^*, and *B2M^KO^* genotypes initiated with Lenti:Cre^BC-sgTS11-IM^, Lenti:mCh/Cre^BC-sgTS11-IM^, or Lenti:mCh-SIIN/Cre^BC-sgTS11-IM^ in *K;Cas9* mice. Tumors were analyzed using the “Adaptive sampling method” (Fig. S6A). **B.** Quantification of relative LN mean tumor size for the 11 tumor suppressor genotypes initiated with Lenti:Cre^BC-sgTS11-IM^, Lenti:mCh/Cre^BC-sgTS11-IM^, or Lenti:mCh-SIIN/Cre^BC-sgTS11-IM^ in *K;Cas9* mice. Tumors were analyzed using the “Adaptive sampling method” (Fig. S6A). **C.** Quadratic fit of the LN mean tumor size between Lenti:Cre^BC-sgTS11-IM^ and Lenti:mCh-SIIN/Cre^BC-sgTS11-IM^ across different sgRNA in the BC-sgTS11-IM pool. Tumors were analyzed using the “Adaptive sampling method” (Fig. S6A). **D.** The relationship between the Relative Resistance metric of mCh-SIIN/Cre^BC-sgTS11-IM^ tumors and the relative LN mean tumor size of Cre^BC-sgTS11-IM^ tumors across different sgRNAs in the BC-sgTS11-IM pool.

**Fig 7.**
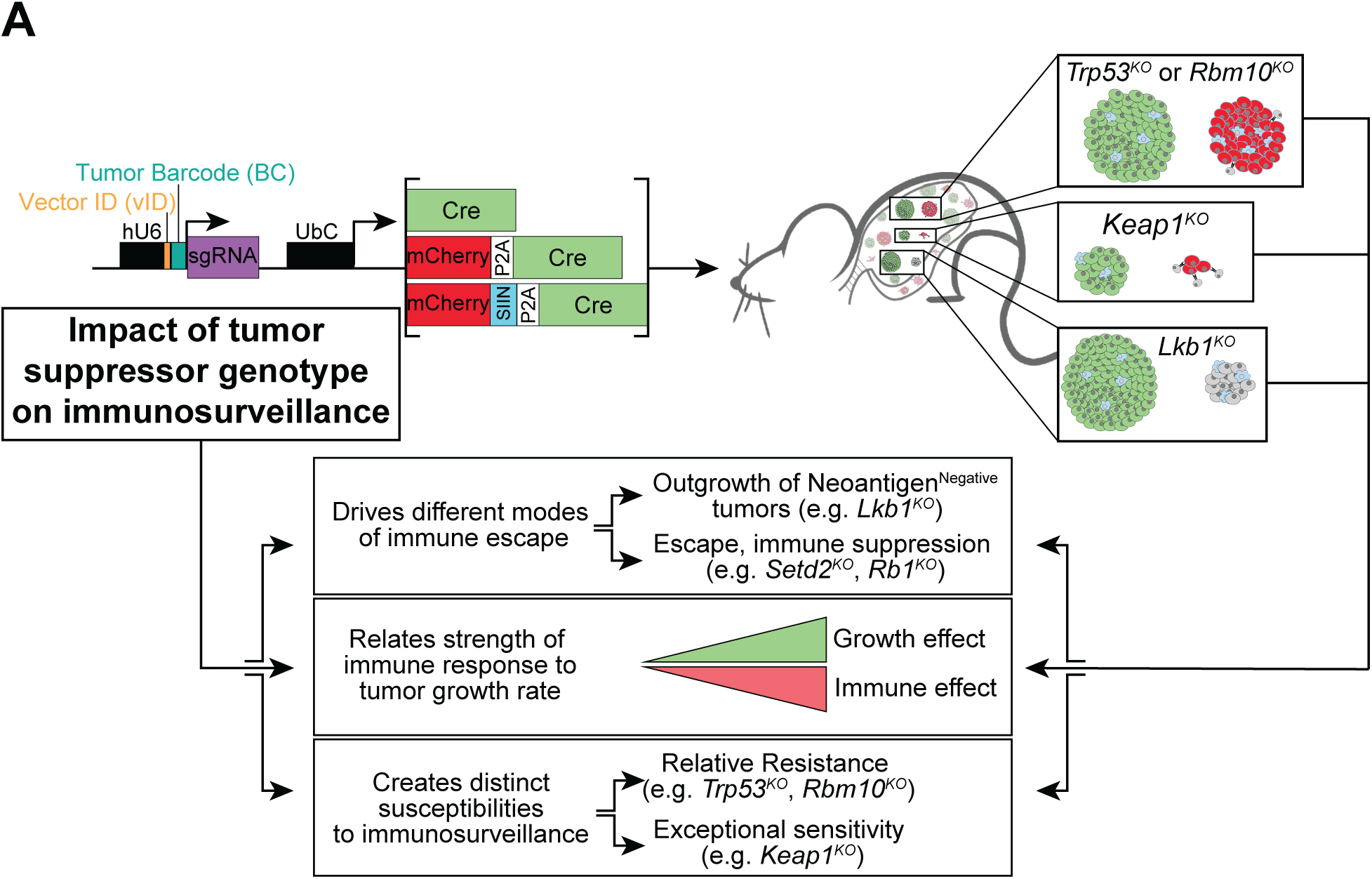
The degree of immunosurveillance is tightly linked to the general rate of tumor expansion with key exceptions for inactivation of specific tumor suppressor genes *Trp53*, *Rbm10*, and *Keap1*. **A.** One-at-a-time gene knockout approaches reveal that tumor suppressor genotype is an important mediator of the immune response mounted against the highly immunogenic SIINFEKL neoantigen. In the context of *Setd2^KO^* and *Rb1^KO^*, tumor suppressor gene inactivation sufficiently promotes immunoevasion such that tumors can grow despite sustained neoantigen expression. However, *Lkb1^KO^* tumors require the selection of neoplastic cells which have silenced neoantigen expression and surface presentation in order to sufficiently evade immune detection. Using a multiplexed high throughput *in vivo* screen combining CRISPR/Cas9-mediated genome editing, neoantigen expression, and tumor barcoding, we simultaneously investigated the impact of 11 tumor suppressor gene inactivations on immunosurveillance in three distinct and increasingly immunogenic contexts. Surprisingly, tumor suppressor genotypes that most strongly promote tumor outgrowth in poorly immunogenic contexts are most sensitive to immunosurveillance in highly immunogenic contexts. However, specific tumor suppressor gene inactivations were exceptions where *Trp53^KO^* and *Rbm10^KO^* promoted resistance to immunosurveillance and *Keap1^KO^* promoting exceptional sensitivity to immunosurveillance.

### Tumor suppressor gene inactivation has complex effects on immune surveillance in immunogenic tumors

To determine the impact of tumor suppressor gene inactivation on immunosurveillance, we quantified metrics of tumorigenesis for each tumor genotype relative to TS^WT^ tumors in each immunogenic condition (Fig S6, Methods). In tumors initiated by Lenti:Cre^BC-sgTS11-IM^, tumor suppressor gene inactivation generated the expected tumor growth-promoting effects (Fig. 6B, Fig. S7, D to F) (*22, 25*). In the context of each tumor suppressor gene inactivation, expression of neoantigens generally had a growth suppressing effect that correlated with the potency of each neoantigenic context. However, *Trp53^KO^, Rbm10^KO^*, and to a lesser extent *Smad4^KO^* tumors initiated by Lenti:mCh-SIIN/Cre^BC-sgTS11-IM^ were similar in relative tumor size to those initiated by Lenti:Cre^BC-sgTS11-IM^, suggesting that inactivation of these genes promotes resistance to tumor immunosurveillance. In contrast, *Keap1^KO^* and *Arid1a^KO^* tumors initiated by Lenti:mCh-SIIN/Cre^BC-sgTS11-IM^ were significantly smaller in relative tumor size to TS^WT^ tumors suggesting that inactivation of these genes further promotes tumor immunosurveillance.

The differential effects of tumor suppressor gene inactivation across immunogenic contexts prompted us to determine if there was a relationship between potency of tumor suppression in poorly immunogenicity contexts and the sensitivity to immune surveillance in highly immunogenic contexts. To determine whether specific genetic contexts strongly affected tumor immunosurveillance, we directly compared the log-normal (LN) mean tumor size in tumors initiated by Lenti:Cre^BC-TS11-IM^ to that of tumors initiated by Lenti:mCh-SIIN/Cre^BC-TS11-IM^. Unexpectedly, tumor genotypes that more strongly promoted tumor outgrowth in poorly immunogenic contexts were significantly more susceptible to immune-mediated clearance in a highly immunogenic context (Fig. 6C, Fig. S8A). This trend was not observed when comparing tumors initiated by Lenti:Cre^BC-TS11-IM^ to tumors initiated by Lenti:mCh/Cre^BC-TS11-IM^, and as such, this proliferation-dependent sensitivity to immunosurveillance appears reliant on a highly immunogenic stimulus (Fig. S8B).

To directly assess the sensitivity of tumors to immune surveillance across genotypes, we calculated a resistance metric (R) by normalizing the relative LN mean tumor size in the highly immunogenic context against the relative LN mean tumor size in the poorly immunogenic context. Instead of a constant R across genotypes, we observed a clear negative linear correlation between resistance and tumor growth in the poorly immunogenic context, which did not occur in the context of tumors initiated by Lenti:mCh/Cre^BC-sgTS11-IM^. suggesting that more proliferative tumors are more sensitive to immune surveillance in highly immunogenic conditions (Fig. 6D and Fig. S8, C to E). Through these analyses, we identified *Keap1^KO^* and *Arid1a^KO^ Trp53^KO^*and *Rbm10^KO^* as outliers (Fig. 6D). That *Keap1^KO^* and *Arid1a^KO^* tumors are positioned below the trend-line supports their increased sensitivity to immunosurveillance, while *Trp53^KO^* and *Rbm10^KO^* positioning above the trendline supports their increased resistance to immunosurveillance.

All together, these findings have uncovered both a general negative correlation between a tumor’s resistance to immunosurveillance and its proliferation rate and also highlighted specific tumor genotypes that shape tumor immunosurveillance during early tumor development (Fig. 7).

## DISCUSSION

In this study, we systematically and quantitatively assess the impact of tumor suppressor genotype on immunosurveillance in an autochthonous cancer model. Employing one-at-a-time gene knockout approaches within the context of tumors that express increasingly immunogenic neoantigens we discovered tumor genotype-dependent heterogeneity regarding the mechanisms of immune evasion between KRAS-driven tumors that had one of three prevalent tumor suppressor genes inactivated (*Lkb1*, *Setd2*, and *Rb1*). Further, we developed a relatively high throughput *in vivo* tumor barcoding strategy to simultaneously assess 14 distinct tumor genotypes across 3 levels of immunogenicity in parallel within individual mice. Unexpectedly, we identified a strong negative relationship between the rate of tumor growth after inactivation of specific tumor suppressor genes and the resistance of tumors to immunosurveillance. Moreover, we identified multiple tumor suppressor genes that either increase or decrease susceptibility to immunosurveillance. Thus, we have created and employed a toolkit that enables the generation of conditional tumor models with predictable levels of immunogenicity across defined genotypes and identified important positive and negative regulators of immunosurveillance.

Our data strongly indicate that a general promoter of tumor immunosurveillance is the rate of a tumor expansion after its initiation. Inactivation of tumor suppressor genes that elicit the strongest growth promoting effects also had the highest degree of immunosurveillance while inactivation of tumor suppressor genes that have weakest effects on tumor expansion after initiation had the lowest degree of immunosurveillance. In fact, the relationship between the degree of immunosurveillance due to expression of strong neoantigens and the tumor expansion rate due to tumor genotype seems to strengthen in a non-linear manner (Fig. 6C). This finding is unexpected and contrasts with the outcome of a study using transplantable tumor cell line models that reported a general positive correlation between tumor suppressor gene inactivation and evasion of immune surveillance (*17*). Most obviously, the distinction between transplantation of established cancer cell lines that were manipulated in vitro and then injected into an autologous mouse recipient versus the in vivo transformation of normal epithelial cells growing in their natural setting but forced to express neoantigens is the likely explanation for the apparent discrepancy. These very different scenarios likely examine distinct aspects of immune surveillance that deserve additional attention. We speculate that our approach may afford insight into the role of immune pressure during the early phases of cancer formation and thus reveal important features of cancer etiology that may inform strategies for cancer interception.

Our study identified *Trp53* and *Rbm10* as genes whose inactivation promoted resistance to immune surveillance. Consistently, multiple studies have indicated that *Trp53* inactivation promotes immune evasion, and that its re-activation in cancer models promotes anti-tumor immunity (*29–34*). However, more work is needed to understand how *Rbm10^KO^* promotes resistance to immunosurveillance in the context of potent neoantigens, as multiple reports indicate that RBM10 deficient tumors stimulate anti-tumor pathways normally associated with immunosurveillance (*16, 35, 36*). In contrast, we also identified *Arid1a* and *Keap1* whose inactivation sensitized tumors to immunosurveillance in the context of strong neoantigen expression. Consistently, *Arid1a* inactivation is associated with anti-tumor immunity and greater progression-free survival in the context of patients on anti-PD-1 therapy (*37, 38*). More recently, *Arid1a* inactivation was found to promote anti-tumor immunity via increased R-loop-derived cytosolic DNA that activates inflammatory signaling pathways (*39*). On the other hand, *Keap1* inactivation, which had the strongest genotype-dependent anti-tumor effect in our model, is curiously associated with resistance to anti-PD-L1 monotherapies but increased susceptibility to combination anti-PD-L1/anti-CTLA4 checkpoint therapies (*14, 19, 40, 41*). Perhaps, the strong anti-tumor response to neoantigen expression in early tumor cells that have *Keap1* inactivation strongly selects for mechanisms that promote immune evasion and thus become recalcitrant to checkpoint monotherapy.

Although this study uncovered an unexpected correlation between the potency of tumor suppressor gene inactivation and sensitivity to immunosurveillance in highly immunogenic contexts, little is currently understood about the mechanism by which this occurs. Furthermore, little is understood about the difference in how tumor suppressor gene inactivations regulate the immune response in highly immunogenic contexts compared to poorly or mildly immunogenic contexts. Mechanistic investigation as to why specific tumor suppressor gene inactivations either promote or limit immunosurveillance of tumors in highly immunogenic contexts may uncover new therapeutic vulnerabilities. In addition, while this study highlights the benefit of mixing vectors of different immunogenicities within a single mouse to uncover tumor suppressor gene inactivations which either promote sensitivity or resistance to immunosurveillance, we found that the presence of highly immunogenic tumors impacted the initiation and outgrowth of poorly immunogenic tumors across all genotypes. Namely, in tumors initiated by the poorly immunogenic Lenti:Cre vector, tumor suppressor gene inactivation decreased tumor initiation compared to TS^WT^ tumors initiated by the same vector. It may prove insightful to determine the effects of homogenizing the degree of immunogenicity across each tumor in a single animal to better understand the extent to which the presence of highly immunogenic tumors impact the immunosurveillance of poorly immunogenic tumors.

Together, these findings highlight the importance of tumor suppressor gene inactivation in the process of immunoediting and show that individual gene inactivations differentially regulate immunosurveillance in response to fixed neoantigens. Immunoediting has typically been thought to begin well after malignant transformation and to rely primarily on the acquisition of passenger mutations to generate neoantigens and randomly promote the selection of tumor cells which better evade immune detection. However, these findings may point to a more significant role for driver mutations in regulating the immune response to neoantigens throughout the process of immunoediting. A similar study focusing on the importance of oncogenic driver mutations on immunoediting of tumors expressing fixed neoantigens would further strengthen our findings that driver mutations can regulate immunoediting. Additionally, gaining mechanistic insight as to how these tumor suppressor gene inactivations regulate immunosurveillance throughout tumor development may provide insight as to why certain mutations promote resistance or sensitivity to immune checkpoint therapies and may uncover therapeutic vulnerabilities that increase the efficacy of these therapies (*12, 13, 15, 18, 38*).

## MATERIALS AND METHODS

### Animal Studies

All work was performed under compliance with Institutional Animal Care and Use Committee at the University of Pennsylvania (#804774). *Kras^LSL-G12D^* (*K*) mice (Jax stock number 008179) and *Rosa26^LSL-Cas9::EGFP^* (*Cas9*) mice (Jax stock number 026175) were maintained on a B6J background. Tumors were transduced via endotracheal inhalation of a lentivirus that expresses Cre-recombinase (*42*). To initially determine how each vector modulates immunoediting, *K* mice were transduced with 1x10^5^ plaque forming units (PFU) per mouse of Lenti:Cre, Lenti:mCh/Cre, or Lenti:mCh-SIIN/Cre. To test how specific TSGs impact immunoediting, *K;Cas9* mice were transduced with 6x10^4^ PFU of Lenti:mCh/Cre or Lenti:mCh-SIIN/Cre containing an inert, non-targeting sgRNA or a sgRNA targeting *Lkb1*, *Setd2*, or *Rb1*. For the Tuba-seq screen, each *K;Cas9* mouse was transduced with a combination of Lenti:Cre^BC-sgTS11-IM^, Lenti:mCh/Cre^BC-sgTS11-IM^, and Lenti:mCh-SIIN/Cre^BC-sgTS11-IM^ that contained the barcoded sgRNA pool (6x10^4^ PFU per virus, 1.8x10^5^ PFU total per mouse). *K* mice in this screen were transduced with the same three viruses, but using 1x10^5^ PFU per virus per mouse (3x10^5^ PFU total per mouse).

### Immunohistochemistry

Lungs were inflated using a 10% neutral-buffered formalin and then placed into the formalin solution overnight at room temperature before dehydration in a graded alcohol series. Paraffin-embedded and H&E sections were produced by the Penn Molecular Pathology and Imaging core. Immunohistochemistry was performed after a citrate-based antigen retrieval using the following antibodies: CD45 (1:100, Biolegend, 103102), CD3 (1:700, Abcam, ab5690), mCherry (1:500, Novus Biologicals, NBP2-25156SS), CD8 (1:500, Cell Signaling, 98941S, clone D4W2Z), FoxP3 (1:500, Cell Signaling, 12653S, clone D6O8R), Ly6G (1:500, Biolegend, 127601, clone IA8), and Arginase I (1:1000, Thermo Scientific, PA5-29645). Primary antibodies were incubated overnight at 4°C. After primary antibody staining, an ABC reagent for rabbit (Vector Laboratories, PK-4001) or rat (Vector Laboratories, PK-6104) and ImmPACT DAB (Vector laboratories, SK-4105) were used per manufacturer’s instructions. Sections were counterstained with Hematoxylin (1:10, Sigma-Aldrich, GHS316-500ML). IHC staining was quantified on ImageJ using Mean Gray Value as previously described (*43*). All mean gray values were then divided by 255 to convert the value to percent positive area.

Individual tumor area and tumor burden were analyzed using ImageJ. Each tumor was individually circled on ImageJ using the freehand selection tool. The area was measured for each tumor in pixels on ImageJ. Tumor burden was calculated by dividing the sum of tumor area within a lung divided by the total lung area.

Qualitative assessments of mCherry were done using 20x IHC images. Each image was assessed and banked into one of three groups: negative, partial, or positive depending on its mCherry expression status. The number of tumors in each category was summed and divided by the total number of tumors assessed to determine percentages for each group. Chi-squared analyses were used to determine statistical significance.

### Plasmids

All vectors used in these experiments were derived from LentiCRISPRv2Cre (Addgene plasmid #82415). The Ef-1α promoter and Cas9 gene were removed from the plasmid by cutting with EcoRI and BamHI restriction enzymes. All restriction digests were run according to protocols found using NEBcloner (New England Biolabs). Then, a Gibson assembly reaction was used to insert the UbC promoter alone or to insert UbC-mCherry to generate Lenti:Cre or Lenti:mCh/Cre. Lenti:mCh-SIIN/Cre was then generated by linearizing Lenti:mCh/Cre plasmid with XhoI and performing a second Gibson reaction to insert the SIINFEKL DNA sequence directly after mCherry. Each new plasmid was transformed and individual colonies were sequenced by Sanger sequencing to confirm successful cloning.

For sgRNA expression from the U6 promoter, it is necessary to maintain a 23 nt gap between the promoter TATA box and the first transcribed G nucleotide. To account for the vID and BC sequence, the hU6 promoter, spacer, and part of the chimeric sgRNA backbone were removed using KpnI and AfeI. A modified insert containing a shortened spacer and a vector ID was inserted into each vector by Gibson assembly. This insert also restored the hU6 and chimeric sgRNA backbone sequences. vID insertion was confirmed by Sanger sequencing transformed colonies.

### Tuba-seq Plasmid Library Generation and Validation and Individual sgRNA Insertion

sgRNA sequences for Tuba-seq were previously validated and published (*22*). Each sgRNA oligo contained the following sequence: 5’-GGAGACCTGCGTCTCACACCHHHHHHHHHHG-sgRNA sequence-GTTTAGAGACGGCTCCGCGC-3’. The 10nt tumor barcodes only consisted of A, C, or T nucleotides to minimize the chance of early initiation of transcription at an upstream G nucleotide. Oligos were ordered separately and then pooled together at 1ng/uL for PCR amplification using NEB Fusion HS Flex Enzyme and Buffer (New England Biolabs, M0535S),10mM dNTPs, and primers (Forward: 5’ GGAGACCTGCGTCTCACACC 3’, Reverse: 5’ GCGCGGAGCCGTCTCTAAAC 3’). PCR conditions were as follows: 98°C for 2 min, 98°C for 8s*, 65°C for 12s*, 72°C for 10s*, 72°C for 5min, then 4°C indefinitely. The starred conditions were repeated for 20 cycles. PCR product was run on a 2% agarose gel and gel purified using the QIAquick Gel Extraction Kit (Qiagen, 28704). Simultaneously, Lenti:Cre, Lenti:mCh/Cre, or Lenti:mCh-SIIN/Cre plasmids were digested with BsmBI-v2 (New England Biolabs, R0739S). The digest was run on a 1% agarose gel and the digested vector was excised and purified from the gel using the QIAquick Gel Extraction Kit.

Both the sgRNA oligo and the digested vector were further purified using an ethanol precipitation. Then, a golden gate reaction was performed by mixing 500ng of digested vector, 100ng sgRNA oligo insert, 10 units Esp3I (Thermo Fisher, ER0452), 3,000 units T7 ligase (Enzymatics, L6020L), 1mM DTT, and 1mM ATP. Samples were cycled between 37°C and 20°C every 5 minutes overnight for 100 cycles. The new plasmid libraries were purified using the MinElute PCR Purification Kit (Qiagen, 28004) and eluted in 10uL of water.

Plasmid libraries were transformed using MegaX DH10B T1^R^ Electrocomp™ Cells (Invitrogen, C640003). 2uL of the golden gate product was mixed with 20uL of electrocompetent cells and then electroporated in a 1mm cuvette using the following pulse conditions: 25µF capacitance, 200Ω resistance, and 2,000V voltage. Transformed cells underwent 1h of recovery and then were transferred to a flask containing 200mL LB for an overnight incubation in a 30°C shaker. The next day, plasmids were isolated using the PureLink™ Expi Endotoxin-Free Maxi Plasmid Purification Kit (Invitrogen, A31217).

After library generation, sgRNA and tumor barcode diversity were determined by PCR amplification of the sgRNA and tumor barcode region of each plasmid library and sequencing using a MiSeq platform (Azenta Life Sciences). The primers were: Forward: 5’ CCGTAACTTGAAAGTATTTCGATTTCTTGGC 3’, Reverse: 5’ CGGTGCCACTTTTTCAAGTTG 3’.

To clone individual sgRNAs into each vector, the same cloning strategy was used. After the Golden Gate cloning reaction, samples were transformed using One Shot™ Stbl3™ Chemically Competent *E. coli* (Invitrogen, C737303). Individual colonies were picked to confirm the presence of the sgRNA. A single colony with a known tumor barcode sequence was then chosen and a bacterial stock was made.

### Cell Lines

All cell lines were grown in DMEM supplemented with GlutaMAX (Invitrogen, 10566016), 10% fetal bovine serum, and gentamicin (Invitrogen, 15750060) at 37°C and 5% CO_2_.

To create ‘spike in’ cells for Tuba-seq, LG1233 cells were infected with Lenti:mCh/Cre-PuroR (derived from Lenti-CRISPRv2Puro—Addgene #98290) that contained a library of GFP-targeting sgRNAs with unique tumor barcode sequences. The GFP sgRNA sequence was: 5’ CAAGCAGAAGAACGGCATCA 3’. A GFP sgRNA was used since it was not included in the sgRNA library for Tuba-seq and could therefore be distinguished as benchmark DNA rather than tumor DNA. Cells were infected in multiple wells of a 6 well plate using a serial dilution of virus in each well to limit the chance of multiple lentiviral particles infecting a single cell. Transduced cells were selected for using a Puromycin selection as described above. Once selection was complete, 1x10^3^ cells from each initial well were plated in a 15cm plate to generate single clones. Each clone was selected and expanded until DNA could be collected using the Qiagen DNeasy Blood & Tissue Kit (Qiagen, 69504) as instructed. The barcode and sgRNA sequence were amplified from the genomic DNA using the following primer set: Forward: 5’ CCGTAACTTGAAAGTATTTCGATTTCTTGGC 3’, Reverse: 5’ CGGTGCCACTTTTTCAAGTTG 3’. PCR amplicons were Sanger sequenced to confirm the presence of the GFP sgRNA and to determine the tumor barcode sequence associated with each clone. 3 benchmark cell lines were established with the following tumor barcode sequences: CTATTAACAA, CCACTTTCCT, and TACATTATTA.

### Flow Cytometry

Lungs were harvested from mice and one lobe per mouse was set aside for flow cytometry. Lung tissue was cut into small pieces and 1mL of digestion buffer was added to each sample. Each 1mL of digestion buffer contained 700uL HBSS media (-Mg^2+^-Ca^2+^), 100uL Trypsin-EDTA (0.25%), 100uL Collagenase IV (Worthington Biochemical Corp., LS004188, from 10mg/mL stock dissolved in HBSS), 0.5U Dispase (Corning, 354235, 5U/mL stock). Samples were shaken at 37°C for 45min at 200rpm to digest the tissue. Tissue was further dissociated using a p200 pipette. 2mL of quench buffer was added to each sample to halt tissue digestion. Each 2mL of Quench buffer contained 200uL fetal bovine serum and 7.5uL DNase (Roche Diagnostics, 10104159001, from 10mg/mL stock dissolved in water) diluted in 1.8mL Leibovitz’s L-15 media (Gibco, 11415064).

All samples were passed through a 45um filter and red blood cells were lysed with 1mL ACK lysis buffer (Gibco, A1049201) for 5min. The lysis reaction was then quenched with 9mL PBS. Samples were then stained a cocktail of primary antibodies and viability dye for in the dark 30 min at 4°C. LIVE/DEAD™ Fixable Near IR (Invitrogen, L34992) was used as directed by the manufacturer. The antibodies used were: CD45 (1:200, eFluor 450, Invitrogen, 48-0451-82, clone 30-F11), H2Kb (1:200, BV510, Biolegend, 116523, clone AF6-88.5), and H2Kb-SIINFEKL (1:100, APC, Biolegend, 141605, clone 25-D1.16).

Cells were then washed twice with FACS buffer. Finally, cells were resuspended in FACS buffer and flow cytometry was performed using an Attune NxT flow cytometer (Thermo Fisher). Data were analyzed in FlowJo. Tumor cells were gated on CD45^Negative^GFP^Positive^ live singlets (Fig. S1A).

### Lentivirus Production

HEK 293FT cells were transfected with lentiviral plasmid, Δ8.2, and VSV-G plasmids in a 4:3:1 ratio diluted into Opti-MEM™ I Reduced Serum Medium (Invitrogen, 31985062). Polyethylenimine was used to increase the transfection efficiency. 24 hr after transfection, media was replaced with fresh DMEM supplemented with 25 mmol/L HEPES (Gibco, 15630-080) and 3 mmol/L caffeine (Sigma, C0750). The next day, virus media was collected from each plate and filtered through a 0.45um PVDF filter (Foxx Life Sciences, 378-3415-OEM). Then, virus media was centrifuged at 107,000xg for 2h at 4°C. The remaining virus media was poured out and virus pellets soaked overnight in 100uL of PBS at 4°C. After this, pellets were triturated and vortexed at 4°C for 15min before being centrifuged for 30s at 16,000xg to remove debris. Lentivirus was aliquoted and stored at -80°C for later use.

Lentiviruses were titered using Green-Go cells, which are derived from 3TZ cells and contain a Cre-dependent GFP reporter (*44*). 2x10^5^ cells were plated in 6-well plates, and 24 hours later lentivirus was added at 10, 1, and 0.1 μL per well. Cells were analyzed by flow cytometry for GFP expression 48 hours following infection, and viral titer was calculated accordingly.

For Tuba-seq, the plasmid sgRNA libraries generated for Lenti:Cre^BC-sgTS11-IM^, Lenti:mCh/Cre^BC-sgTS11-IM^, and Lenti:mCh-SIIN/Cre^BC-sgTS11-IM^ were individually made into lentivirus. The three viruses were titered separately and only mixed at the time of endotracheal transduction.

### Genomic DNA Extraction for Tuba-seq

Lung lobes were harvested from mice and weighed. Then, each lung was placed into 10mL of digestion buffer containing 100 mM NaCl, 20 mM Tris, 10 mM EDTA, 0.5% SDS, and 100uL of 20mg/mL Proteinase K as previously described (*22*). 5x10^5^ cells from each of the three benchmark cell lines was added to each sample. Tissue was cut into small pieces and incubated at 55°C overnight. Once tissue was digested the next day, DNA was extracted from 5mL of each digested sample by phenol-chloroform extraction. DNA yield and quality was determined using a NanoDrop One^C^ (Thermo Fisher).

### Library Preparation for Sequencing

Library preparation occurred in two steps using a nested PCR strategy. First, the vID, BC and sgRNA sequences were amplified by PCR. Then, Illumina adapter sequences were added to each product in a second, shorter round of PCR.

To amplify the tumor barcode and sgRNA sequence in the first round of PCR, 32ug of gDNA was used as template, split over 8 reactions with a final volume of 100uL each. Q5 Hot Start High-Fidelity 2x Master Mix (New England Biolabs, M0494X) was mixed with gDNA and a universal forward and reverse primer set (Forward: 5’ CGTGACGTAGAAAGTAATAATTTCTTGGGTAG 3’, Reverse: 5’ GAGGCCGAATTCAAAAAAGCACC 3’). PCR conditions were as follows: 94°C for 2 min, 94°C for 30s*, 57°C for 30s*, 68°C for 30s*, 72°C for 7min, then 4°C indefinitely. Starred conditions were repeated for 32 cycles. A small volume of the final product was run on a 2% agarose gel to confirm barcode and sgRNA amplification. Then, a double size selection was performed on the remaining PCR product to remove gDNA and primer dimer using AmPure XP beads (Beckman Coulter, A63881).

Next, a second PCR was used to add unique dual-indexed Illumina adapters to each mouse sample for sequencing. The unique dual-indexed primers contained the following general sequence: Forward: 5’-AATGATACGGCGACCACCGAGATCTACAC-8nt i5 index-ACACTCTTTCCCTACACGACGCTCTTCCGATCT-7-9 random nucleotides-CCGTAACTTGAAAGTATTTCG-3’, Reverse: 5’-CAAGCAGAAGACGGCATACGAGAT-reverse complement of 8nt i7 index-GTGACTGGAGTTCAGACGTGTGCTCTTCCGATCT-7-9 random nucleotides-CGGTGCCACTTTTTCAAGTTG-3’. Illumina TruSeq i5 and i7 index sequences were used and each mouse received a unique combination of both index sequences. Each forward and reverse primer included 7-9 random nucleotides (indicated as N) directly upstream of the plasmid-binding region of the primer to counteract the limited nucleotide diversity in the amplicon library during sequencing, since no PhiX spike-in was used. To set up the PCR for each mouse in the experiment, Q5 Hot Start High-Fidelity 2x Master Mix was mixed with a unique indexed forward and reverse primer set and the size-selected amplicon from the previous PCR. The final reaction volume per tube was 100uL. PCR conditions were as follows: 94°C for 2 min, 94°C for 30s*, 56°C for 30s*, 68°C for 30s*, 72°C for 7min, then 4°C indefinitely. The starred conditions were repeated for 5 cycles to ensure Illumina adapter sequences were added without further amplifying the original PCR product. Reactions from each mouse were pooled into a single tube and 100uL of the pool was run on a 3% agarose gel. The band with the correct fragment length was cut from the gel and gel extracted using the Qiagen MinElute Gel Extraction Kit (Qiagen, 28604). Samples from each mouse were then pooled together and contaminants were removed from the pooled library using an ethanol precipitation. The final concentration and purity of the library was read on a NanoDrop One^C^ (Thermo Fisher) and samples were submitted for sequencing (Azenta Life Sciences). The pooled sample was sequenced on the Illumina HiSeq platform using a sequencing read length of 2x150bp.

### Raw reads parsing and cleaning

Paired-end reads are first merged using AdapterRemoval (*45*), and merged reads are parsed using regular expressions to identify the 4-nucleotide vID, 10-nucleotide BC and 20-nucleotide sgRNA sequence. For identifying sgRNA sequences, we required a perfect match with the designed sequences, as a mutated sgRNA could cause reduced efficiency or off-target effects. When we encountered low-frequency clonal barcodes within a 1-hamming distance of high-frequency clonal barcodes, we attributed them to sequencing or PCR errors (spurious reads). These low-frequency barcodes were merged with barcodes of higher frequencies. Finally, total reads for each vID-BC-sgRNA combination are calculated.

### Clonal tumor size estimation

Read counts associated with clonal tumors are converted into absolute neoplastic cell numbers by normalizing the BC read counts of tumor cells to the BC reads of the ‘spike-in’ cells added just before tissue digestion. For each sample, we check the read ratio among the three ‘spike-in’ cell lines. If one ‘spike-in’ cell line is under-amplified or over-sampled compared to the other two, we only used the other two to calculate the cell number conversion. Otherwise, all three ‘spike-in’ cell lines were used for the conversion. Samples were sequenced sufficiently deep such that across all experiments we achieved at least 15 cells per read. To perform statistical comparisons of tumor genotypes, we imposed a minimum tumor size cutoff of 100 cells to distinguish potential non-expansion cells, spurious tumors from sequencing errors and truly expanding tumors.

### Plasmid library quality control

To ensure equal representation of sgRNAs across Lenti:Cre^BC-sgTS11-IM^, Lenti:mCh/Cre^BC-sgTS11-IM^, and Lenti:mCh-SIIN/Cre^BC-sgTS11-IM^ libraries, we sequenced the plasmid libraries and confirmed that the read fractions for individual sgRNAs correlated highly across all three vectors (Pearson’s r ≥ 0.99, Fig. S9, A to C). Furthermore, we validated that the sgRNA representation in tumors from *K* mice exhibited a strong correlation with plasmid representation of each genotype (Pearson’s r ≥ 0.95, Fig. S9, D to F). Additionally, we ensured that all designed sgRNAs were present in the library and that the number of clonal barcodes associated with each sgRNA was sufficiently large.

### Filtering low quality samples

First, we removed one K mouse where Lenti:mCh-SIIN/Cre^BC-sgTS11-IM^ did not reduce tumor number or area as observed across other *K* and *K;Cas9* mice.

Then, we identified potential contamination during sample preparation by evaluating the overlap of Lenti:Cre^BC-sgTS11-IM^ tumors with shared vID-BC-sgRNA combinations (Figure S9G). A specific vID-BC-sgRNA combination appearing in more than one sample can either be attributed to random sampling events, as all tumors originate from the same pool of lentiviral particles, or to contamination between samples. If the overlap of vID-BC-sgRNA combinations is due to random sampling, the number of tumors with the same vID-BC-sgRNA combinations appearing in both samples should be predictable as a function of the total number of tumors in each sample. Therefore, we used the total tumor counts from both samples to predict the expected number of tumors with shared vID-BC-sgRNA combinations (Fig. S9H). Upon applying this approach, we identified a sample pair—sample 29791 and sample 29837—with a significantly higher number of shared vID-BC-sgRNA combinations than expected. Further investigation into the tumor sizes within these samples revealed two distinct groups: one that aligned well with the predicted overlap due to random sampling, and another indicative of contamination, specifically from sample 29791 to sample 29837 (Fig. S9I). Considering that approximately 50% of the tumors in sample 29837 exhibited shared vID-barcode-sgRNAs, we concluded that sample 29837 was severely contaminated by sample 29791. Due to the extent of the contamination, we decided to exclude sample 29837 from further analysis.

Finally, we identified samples in which gene knockouts had unexpected effects using principal component analysis (PCA) plots using both the relative geometric mean of tumor size and relative tumor counts as metrics. We identified one sample with a distinct pattern across both metrics and therefore excluded this sample from further analysis.

### Summary statistics for overall growth rate

To quantify the impact of each gene on tumor growth, we used three key measures: the size of tumors at defined percentiles of the distribution (specifically the 50^th^, 70^th^, 80^th^, 90^th^, and 95^th^ percentile tumor sizes), the log-normal mean (LN mean) size and geometric mean size (*24, 46*). To standardize the gene knockout effect across different immune contexts, we employed adaptive sampling (*47*). Using tumors initiated by Lenti:Cre^BC-sgTS11-IM^ as the reference, this approach ensured an equal number of tumors were sampled per infectious unit of virus delivered for each sgRNA in each immune context, facilitating consistent calculation of tumor metrics.

### Summary statistics for tumor number and tumor burden

In addition to the tumor size metrics, we characterized the effects of gene inactivation on tumorigenesis by evaluating the number of tumors (“tumor number”) and total neoplastic cell number (“tumor burden”) associated with each genotype. Unlike tumor size metrics, tumor number and burden are linearly influenced by lentiviral titer and sensitive to differences in the representation of each immunogenic vector in the viral pool. As such, to assess the extent to which a given gene (*X)* affects tumor number, we therefore normalized the number of sg*X* tumors in *K;Cas9* mice by the number of sg*X* tumors in the *K* mice:

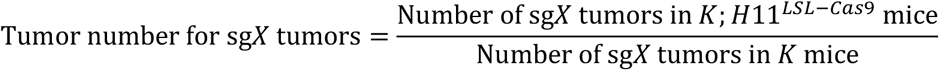

Analogous to the calculation of relative tumor number, we characterized the effect of each gene on tumor burden by first normalizing the sgX tumor burden in *K;Cas9* mice to the burden in *K* mice:

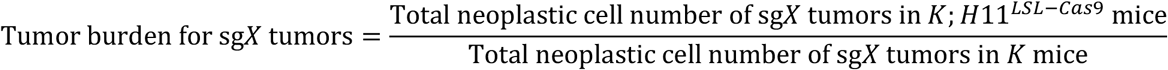

### Model for genetic and immune effect on tumor metrics

We used a mathematical model to dissect the contributions of genetic context, immune response, and genotype-specific immune response to tumor fitness:

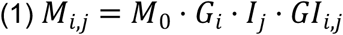

*M_i,j_* is a tumor metric for tumor targeted by sgRNA *i* (refer as genotype *i* hereafter) under the immune context *j*. sg*Inert* (*i = Inert*) is the reference genotype and the immune context faced by the tumor initiated by Lenti:Cre^BC-sgTS11-IM^ (Control tumors) is the reference immune context. *M_0_* is the baseline tumor metric for sg*Inert* tumor under the control context. *G_i_* represents the genetic effect of genotype *i* on tumor fitness. *G_Inert_* =1 and a *G_i_* > 1 indicate a tumor suppressor effect. *I_j_* represents the effect of immune context *j* on tumor fitness. *I_Control_* = 1 and a *I_j_* < 1 indicates the immune context *j* suppresses tumor growth more than the control immune context. *GI_i,j_* captures the interaction between the genotype *i* and the immune context *j* and indicates whether the genotype *i* confers resistance or susceptibility to the immune pressure*. GI_Inert,j_* = *GI_i,Control_* = 1. A *GI_i,j_* >1 indicates the focal genotype *i* is more resistant to the immune response than sg*Inert* tumor under the immune context *j*.

To measure the genetic effect of gene knockout (*G_i_*), we calculated the relative tumor metrics under the control immune context (*RM_i,j=Control_*):

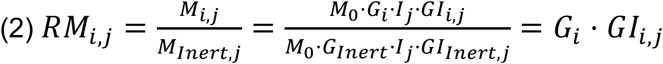

When *j* = *Control*, Equation (2) becomes:

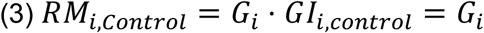

To measure the effect of the immune context (*I_j_*), we calculated the resistance (*R_Inert,j_*) of tumors to a specific immune context j by dividing a tumor metrics of sgInert tumors under the context *j* by the corresponding metrics under the immune control context:

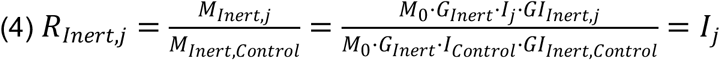

To measure the genotype-specific response of genotype *i* under the immune context *j,* we calculated the relative resistance (*RR*) by diving the *RM* under the immune context *j* (Equation (2)) by the *RM* under the control immune context (Equation (3)):

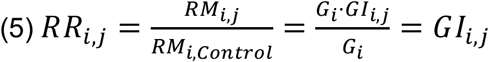

### Calculation of confidence intervals and *p*-values for tumor growth and number metrics

Confidence intervals and *p*-values were calculated using bootstrap resampling to capture variability both between and within mice. We used a two-step nested bootstrap: first resampling mice, then resampling tumors within each mouse, repeating this process 10,000 times. The 95% confidence intervals were derived from the 2.5th and 97.5th percentiles of the bootstrapped values.

The null hypothesis assumes no genotype effect on tumor growth, implying a test statistic equal to 1. Two-sided *p*-values were calculated as:

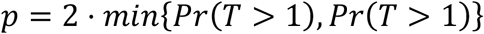

where *T* represents the test statistic and probabilities were empirically estimated from bootstrap resampling. To adjust for multiple testing, *p*-values were corrected using the Benjamini-Hochberg procedure (*48*).

### Fitting linear and quadratic relationships between tumor sizes of different immune context

To explore the relationship between tumor sizes under different immune contexts for the same genotype, we performed linear and quadratic fittings using the LN mean tumor size data. All genotypes, except *B2M* and *PD-L1*, were initially included in the analysis. Tumor sizes were log-transformed to normalize distributions and stabilize variance.

We began by fitting a quadratic model to the relationship between the LN mean tumor sizes of mCh-SIIN/Cre^BC-sgTS11-IM^ tumors and Cre^BC-sgTS11-IM^ tumors. To account for potential outliers, the RANSAC (Random Sample Consensus) algorithm was applied during the fitting process. This robust approach identified *Keap1* as an extreme outlier, which was subsequently excluded from the dataset. *Keap1*’s extreme deviation was interpreted as an outlier that could significantly distort the model.

Following the exclusion of *Keap1*, both linear and quadratic models were refitted to the remaining data. The performance of these models was evaluated using the coefficient of determination (R²) and residual analysis to assess goodness-of-fit and determine the appropriateness of each model.

## Acknowledgements

We would like to thank ULAR staff for animal husbandry, the Molecular Pathology and Imaging Core (MPIC) for histological analysis, Chengcheng Jin for giving us *Cas9* mice, as well as Junwei Shi and Qingzhou Chen for advice on CRISPR library generation and sequencing.

## Author contributions

Conceptualization: KMA, HX, MMW, DMF

Methodology: KMA, HX, MMW, DMF

Formal analysis: KMA, HX

Investigation: KMA, ACG, VIN, MR, KRD

Writing—original draft: KMA, HX, MMW, DMF

Visualization: KMA, HX, MMW, DMF

Supervision: DAP, MMW, DMF

Funding acquisition: DAP, MMW, DMF

## Data and materials availability

The sequencing data set generated and analyzed during the current study will be submitted to the Gene Expression Omnibus database. Other data and relevant code will be available in GitHub.

---

**Fig. S1:**
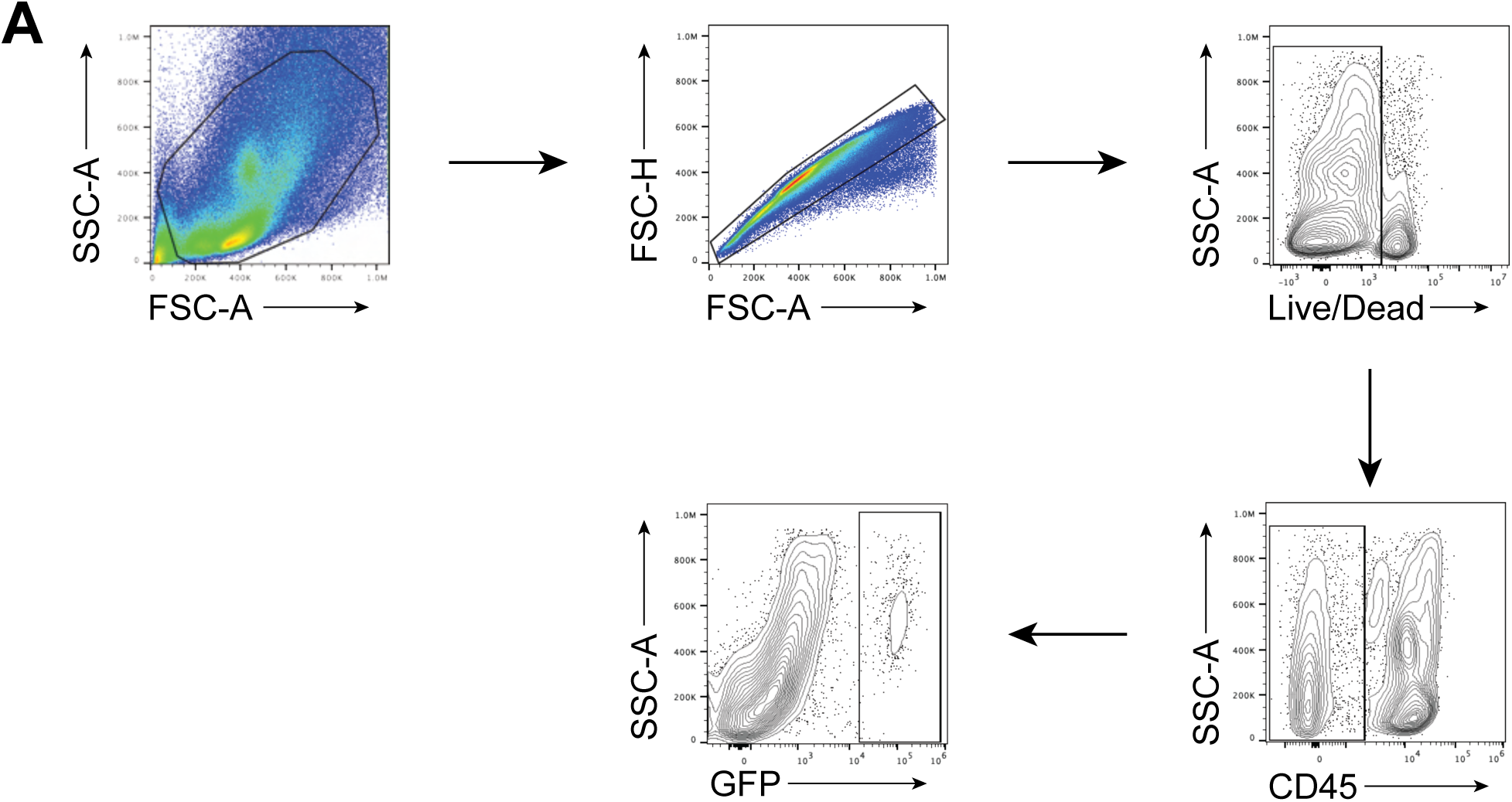
Gating scheme for *in vivo* analysis of neoplastic cells. **A.** Gating strategy for flow cytometry to analyze GFP^Positive^ neoplastic cells.

**Fig. S2:**
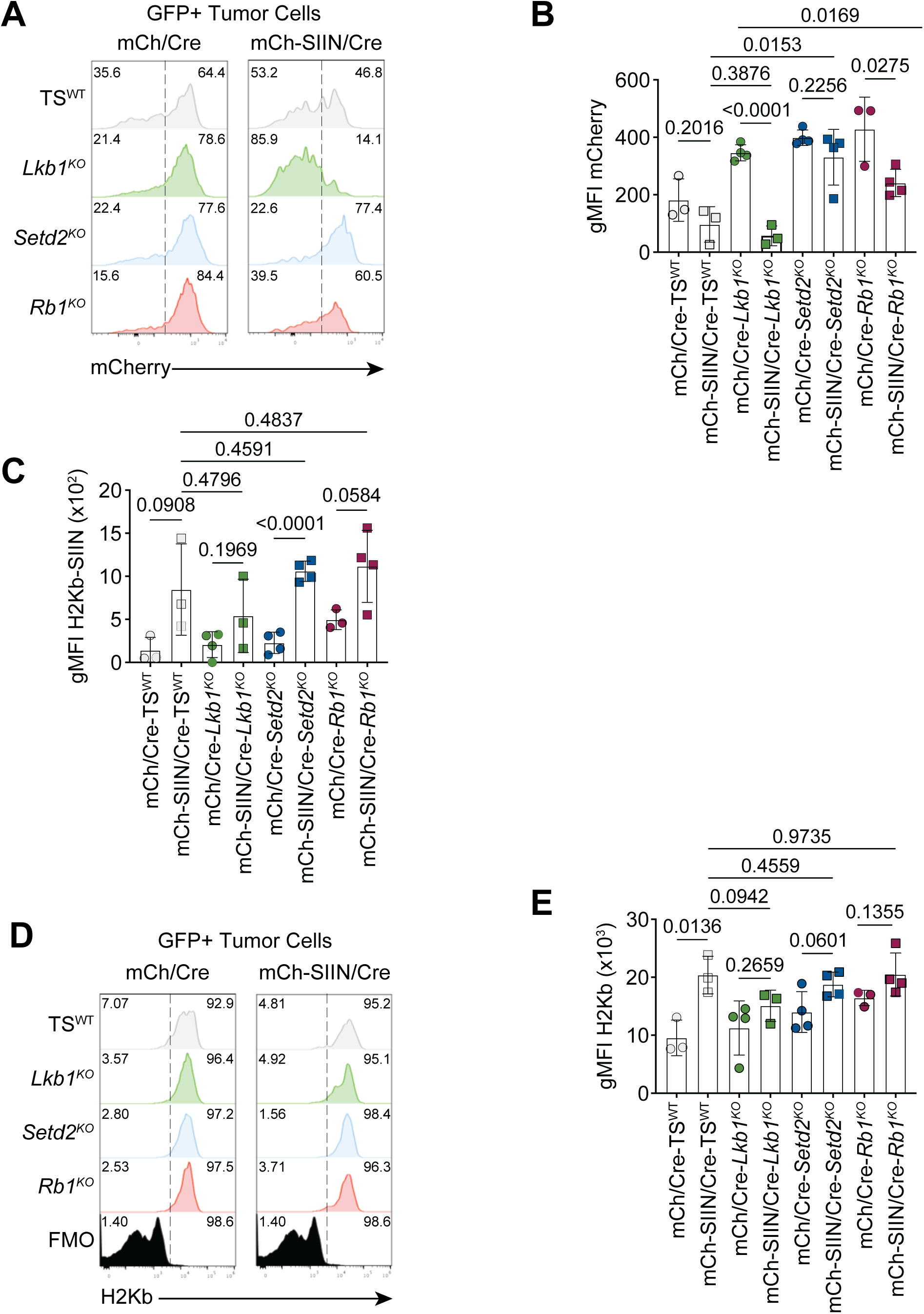
Tumors with *Setd2^KO^* or *Rb1^KO^* maintain mCherry expression and antigen presentation in a highly immunogenic context. **A.** Histogram plots showing mCherry expression measured by flow cytometry across each genotype specified. Plots are separated based on initiating vector and are normalized to mode. Dotted line separates positive and negative populations and value in the upper corner indicates the percentage of cells in that population. **B.** Quantification of gMFI of mCherry expression as measured by flow cytometry, corresponding to the flow plots shown in Supplementary Figure 2A. Statistical significance was determined using unpaired Student’s *t*-tests. Error bars represent mean ± standard deviation. n=4 for all experimental groups except for mCh/Cre-TS^WT^, mCh/Cre-*Rb1^KO^*, mCh-SIIN/Cre-TS^WT^, and mCh-SIIN/Cre-*Lkb1^KO^* (n=3). **C.** Quantification of gMFI of H2Kb-SIINFEKL presentation as measured by flow cytometry, corresponding to the flow plots shown in Figure 3G. Statistical significance was determined using unpaired Student’s *t*-tests. Error bars represent mean ± standard deviation. n=4 for all experimental groups except for mCh/Cre-TS^WT^, mCh/Cre-*Rb1^KO^*, mCh-SIIN/Cre-TS^WT^, and mCh-SIIN/Cre-*Lkb1^KO^* (n=3). **D.** Histograms showing H2Kb presentation on tumor cells measured by flow cytometry across each genotype specified. Plots are separated based on initiating vector and normalized to mode. Dotted line separates positive and negative populations and value in the upper corner indicates the percentage of cells in that population. **E.** gMFI of H2Kb presentation as measured by flow cytometry. Statistical significance was determined using unpaired Student’s *t*-tests. Error bars represent mean ± standard deviation. n=4 for all experimental groups except for mCh/Cre-TS^WT^, mCh/Cre-*Rb1^KO^*, mCh-SIIN/Cre-TS^WT^, and mCh-SIIN/Cre-*Lkb1^KO^* (n=3).

**Fig. S3:**
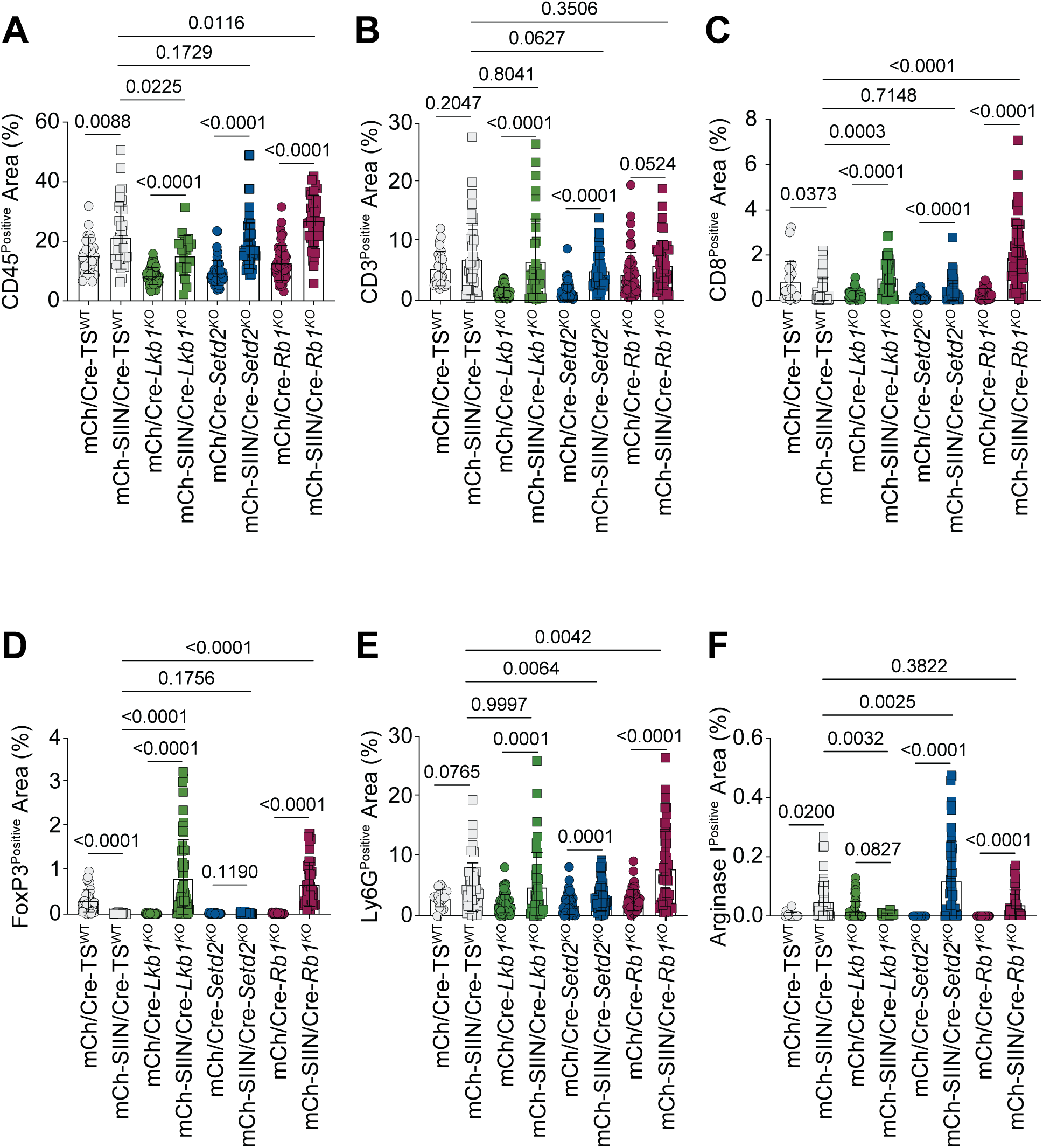
SIINFEKL promotes immune infiltration across tumor suppressor genotypes. **A.** Quantification of CD45 IHC shown in Figure 4A using percent positive area. Statistical significance was determined using unpaired Student’s *t*-tests. Error bars represent mean ± standard deviation. For mCh/Cre-TS^WT^, n=28 tumors from n=4 mice. For For mCh-SIIN/Cre-TS^WT^, n=37 tumors from n=4 mice. For For mCh/Cre-*Lkb1^KO^*, n= 76 tumors from n=4 mice. For mCh-SIIN/Cre-*Lkb1^KO^*, n=20 tumors from n=4 mice. For mCh/Cre-*Setd2^KO^*, n= 45 tumors from n=4 mice. For mCh-SIIN/Cre-*Setd2^KO^*, n=45 tumors from n=4 mice. For mCh/Cre-*Rb1^KO^*, n= 49 tumors from n=3 mice. For mCh-SIIN/Cre-*Rb1^KO^*, n=42 tumors from n=4 mice. **B.** Quantification of CD3 IHC shown in Figure 4A using percent positive area. Statistical significance was determined using unpaired Student’s *t*-tests. Error bars represent mean ± standard deviation. For mCh/Cre-TS^WT^, n=27 tumors from n=4 mice. For mCh-SIIN/Cre-TS^WT^, n=45 tumors from n=4 mice. For mCh/Cre-*Lkb1^KO^*, n= 73 tumors from n=4 mice. For mCh-SIIN/Cre-*Lkb1^KO^*, n=34 tumors from n=4 mice. For mCh/Cre-*Setd2^KO^*, n= 65 tumors from n=4 mice. For mCh-SIIN/Cre-*Setd2^KO^*, n=42 tumors from n=4 mice. For mCh/Cre-*Rb1^KO^*, n= 52 tumors from n=3 mice. For mCh-SIIN/Cre-*Rb1^KO^*, n=45 tumors from n=4 mice. **C.** Quantification of CD8 IHC shown in Figure 4A using percent positive area. Statistical significance was determined using unpaired Student’s *t*-tests. Error bars represent mean ± standard deviation. For mCh/Cre-TS^WT^, n=21 tumors from n=4 mice. For mCh-SIIN/Cre-TS^WT^, n=50 tumors from n=4 mice. For mCh/Cre-*Lkb1^KO^*, n= 72 tumors from n=4 mice. For mCh-SIIN/Cre-*Lkb1^KO^*, n= 37 tumors from n=4 mice. For mCh/Cre-*Setd2^KO^*, n= 68 tumors from n=4 mice. For mCh-SIIN/Cre-*Setd2^KO^*, n=65 tumors from n=4 mice. For mCh/Cre-*Rb1^KO^*, n= 38 tumors from n=3 mice. For mCh-SIIN/Cre-*Rb1^KO^*, n=59 tumors from n=4 mice. **D.** Quantification of FoxP3 IHC shown in Figure 4A using percent positive area. Statistical significance was determined using unpaired Student’s *t*-tests. Error bars represent mean ± standard deviation. ROUT method was used to identify and remove outliers. A Q value of 1 was used for this analysis. For mCh/Cre-TS^WT^, n=36 tumors from n=4 mice. For mCh-SIIN/Cre-TS^WT^, n=53 tumors from n=4 mice. For mCh/Cre-*Lkb1^KO^*, n= 67 tumors from n=4 mice. For mCh-SIIN/Cre-*Lkb1^KO^*, n= 63 tumors from n=4 mice. For mCh/Cre-*Setd2^KO^*, n= 57 tumors from n=4 mice. For mCh-SIIN/Cre-*Setd2^KO^*, n=49 tumors from n=4 mice. For mCh/Cre-*Rb1^KO^*, n= 52 tumors from n=3 mice. For mCh-SIIN/Cre-*Rb1^KO^*, n=54 tumors from n=4 mice. **E.** Quantification of Ly6G IHC shown in Figure 4A using percent positive area. Statistical significance was determined using unpaired Student’s *t*-tests. Error bars represent mean ± standard deviation. For mCh/Cre-TS^WT^, n=17 tumors from n=4 mice. For mCh-SIIN/Cre-TS^WT^, n=49 tumors from n=4 mice. For mCh/Cre-*Lkb1^KO^*, n= 81 tumors from n=4 mice. For mCh-SIIN/Cre-*Lkb1^KO^*, n=42 tumors from n=4 mice. For mCh/Cre-*Setd2^KO^*, n=68 tumors from n=4 mice. For mCh-SIIN/Cre-*Setd2^KO^*, n=65 tumors from n=4 mice. For mCh/Cre-*Rb1^KO^*, n= 53 tumors from n=3 mice. For mCh-SIIN/Cre-*Rb1^KO^*, n=55 tumors from n=4 mice. **F.** Quantification of Arginase I IHC shown in Figure 4A using percent positive area. Statistical significance was determined using unpaired Student’s *t*-tests. Error bars represent mean ± standard deviation. ROUT method was used to identify and remove outliers. A Q value of 1 was used for this analysis. For mCh/Cre-TS^WT^, n=20 tumors from n=4 mice. For mCh-SIIN/Cre-TS^WT^, n=52 tumors from n=4 mice. For mCh/Cre-*Lkb1^KO^*, n= 78 tumors from n=4 mice. For mCh-SIIN/Cre-*Lkb1^KO^*, n=36 tumors from n=4 mice. For mCh/Cre-*Setd2^KO^*, n=68 tumors from n=4 mice. For mCh-SIIN/Cre-*Setd2^KO^*, n=67 tumors from n=4 mice. For mCh/Cre-*Rb1^KO^*, n=47 tumors from n=3 mice. For mCh-SIIN/Cre-*Rb1^KO^*, n=51 tumors from n=4 mice.

**Fig. S4:**
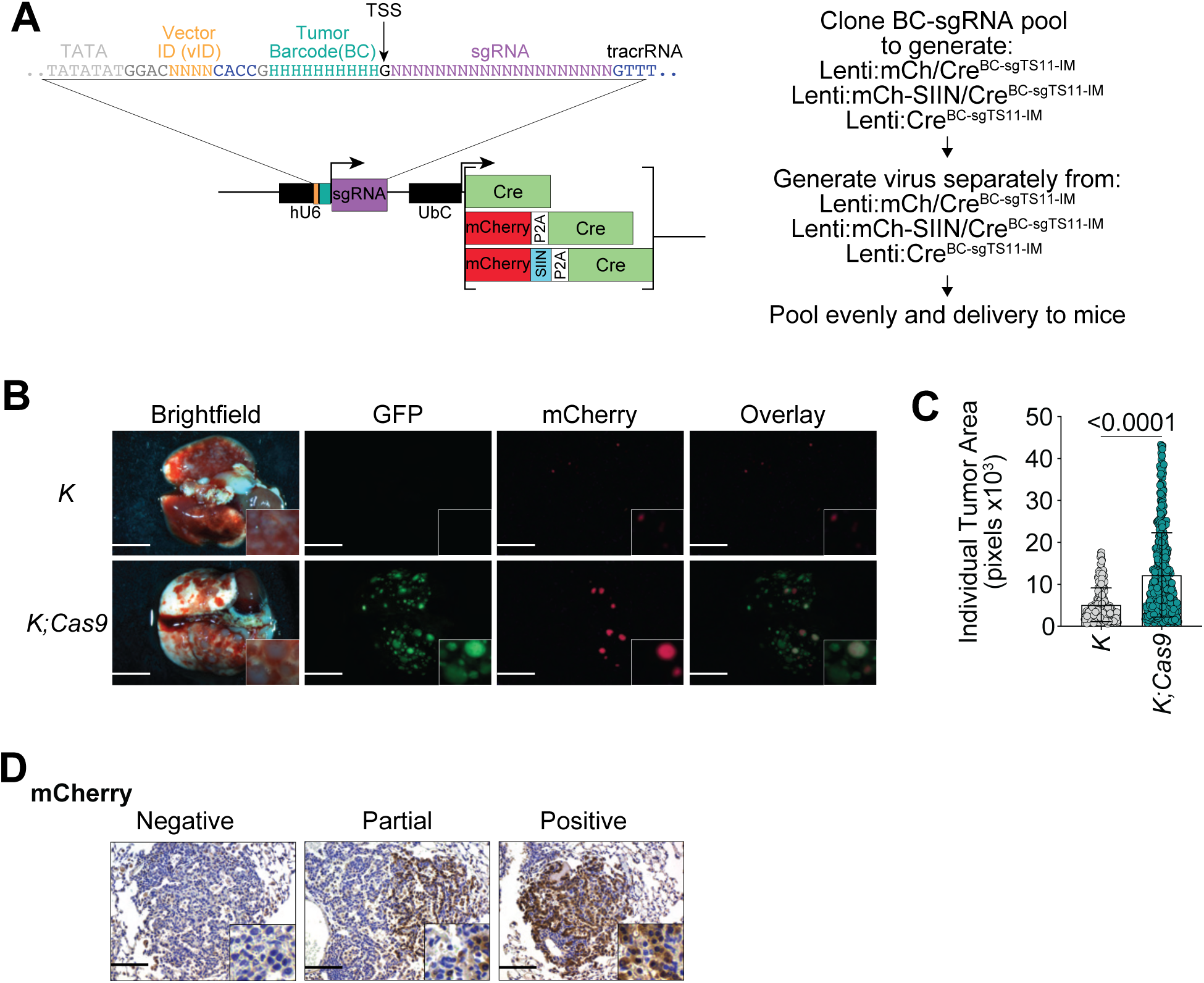
The BC-sgTS11-IM pool sufficiently promotes tumor outgrowth and a subset of tumors are mCherry^Positive^. **A.** Outline of strategy to generate lentivirus pools for Tuba-seq. **B.** Bright field, GFP, and mCherry images of whole dissected lungs. An overlay of the GFP and mCherry images is shown on the far right. Scale bar is 4.4mm. **C.** Quantification of individual tumor area in *K* vs *K;Cas9* mice. Tumor area measured in pixels. Statistical significance was determined using an unpaired Student’s *t*-test. Error bars represent mean ± standard deviation. For *K* mice, n=258 tumors from n=3 mice. For *K;Cas9* mice, n=628 tumors from n=5 mice. ROUT method was used to identify and remove outliers. A Q value of 1 was used for this analysis. **D.** IHC for mCherry in *K* and *K;Cas9* mice. 20x Representative images of different mCherry expression statuses: negative, partial, or positive. Insets are 3x magnified. Scale bar is 119um.

**Fig. S5:**
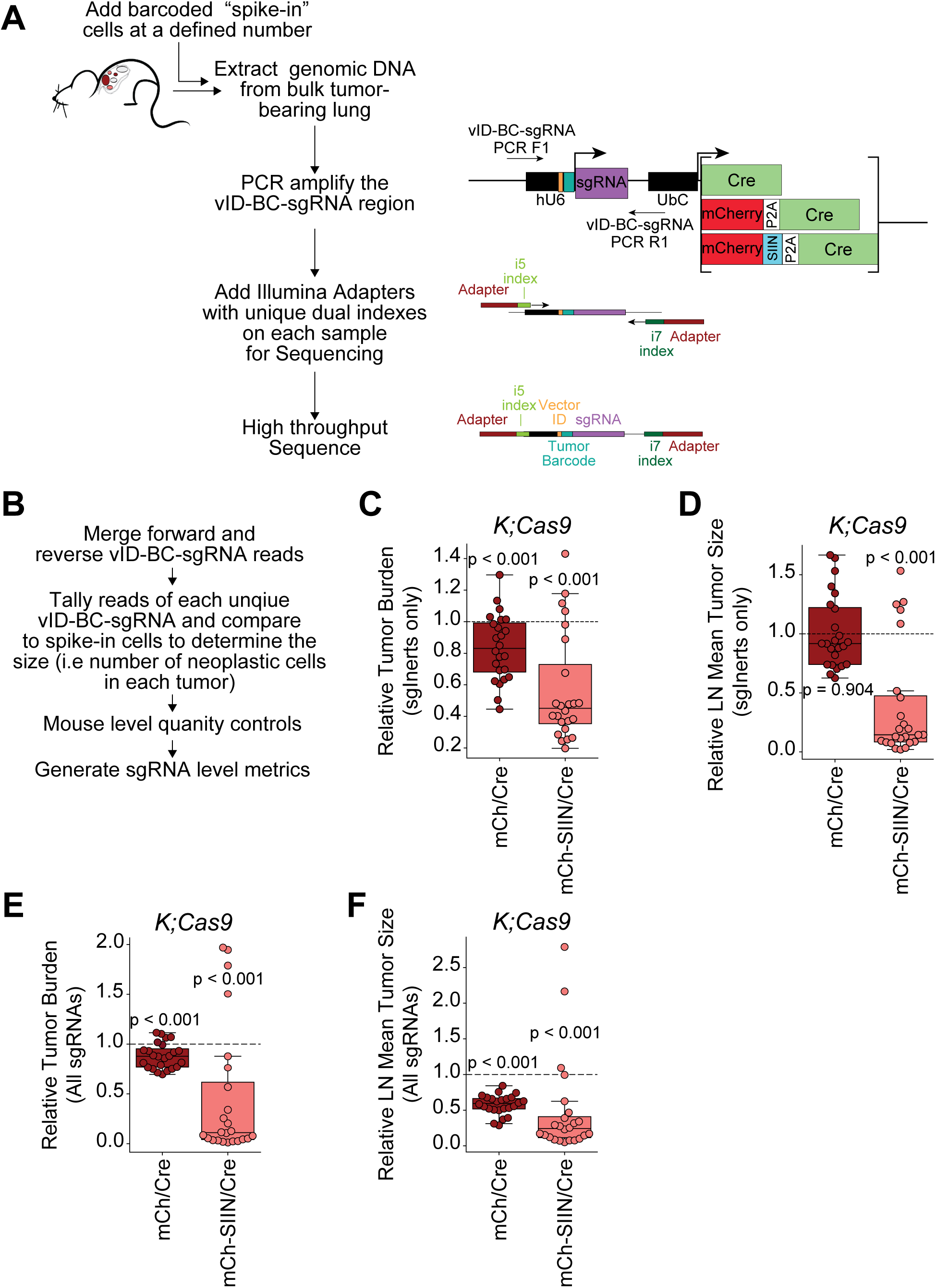
Tuba-seq library preparation and sequencing analysis reveals increasing neoantigen potency restricts tumor outgrowth. **A.** Schematic of nested PCR strategy used for Tuba-seq library preparation before next-generation sequencing. **B.** General summary of steps involved in Tuba-seq analysis after next-generation sequencing. See Methods for more details. **C and D.** Tumor burden (E) or LN mean tumor size (F) of mCh/Cre^BC-sgTS11-IM^ and mCh-SIIN/Cre^BC-sgTS11-IM^ tumors relative to Cre^BC-sgTS11-IM^ tumors across *K;Cas9* mice, focusing exclusively on TS^WT^ tumors. Tumors were analyzed using the “Normal method” (Fig. S6A). One-sample t-tests were used to determine if the ratios were significantly different from 1. **E and F.** Tumor burden (E) or LN mean tumor size (F) of mCh/Cre^BC-sgTS11-IM^ and mCh-SIIN/Cre^BC-sgTS11-IM^ tumors relative to Cre^BC-sgTS11-IM^ tumors in *K;Cas9* mice looking at all sgRNAs except for those targeting *PD-L1* and *B2M*. Tumors were analyzed using the “Normal method” (Fig. S6A). One-sample t-tests were used to determine if the ratios were significantly different from 1.

**Fig. S6:**
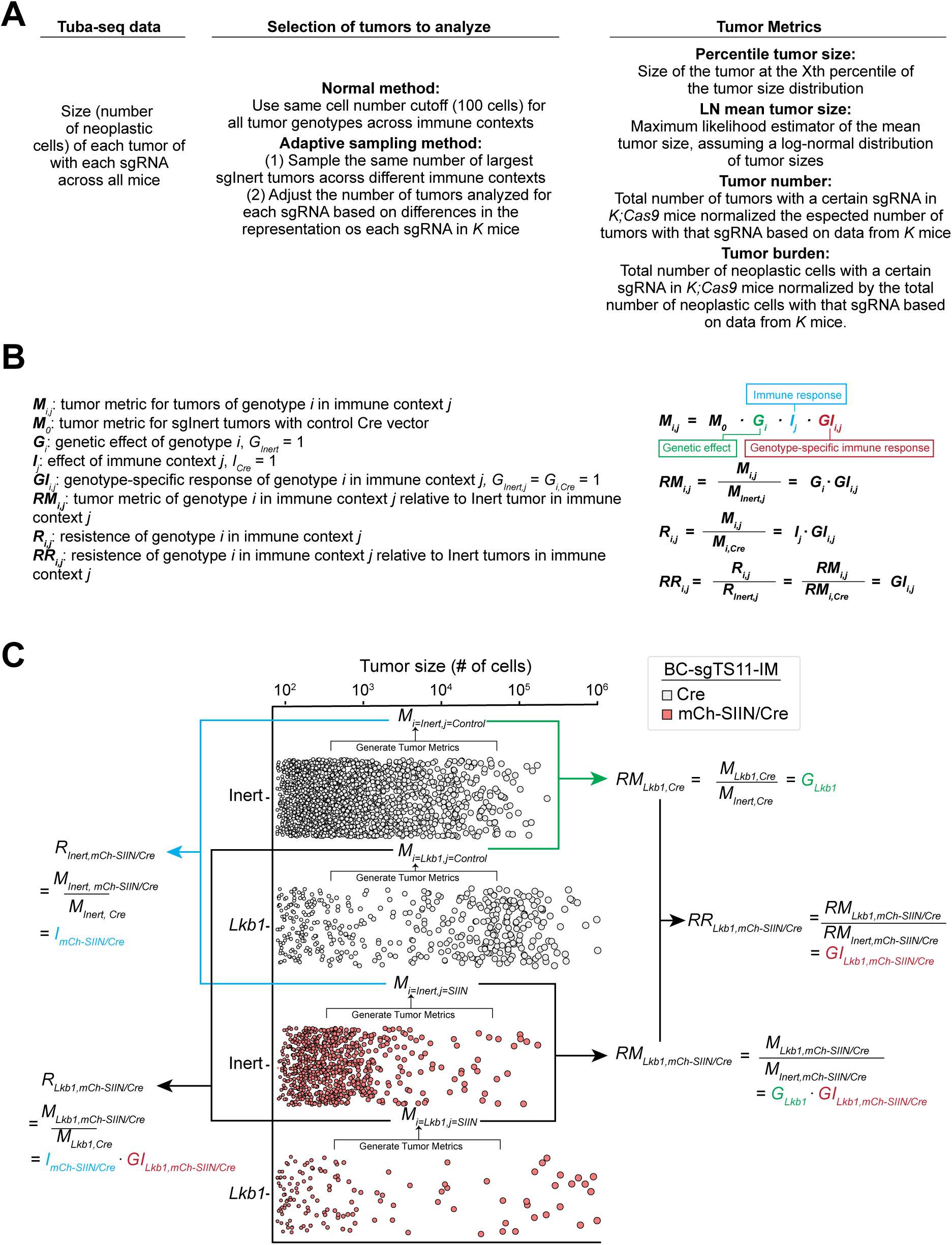
Model and methodology for disentangling tumor fitness contribution. **A.** Explanation of methods and metrics used to analyze Tuba-seq data. **B.** Mathematical explanation of analysis methods used to distinguish the effects of tumor suppressor gene inactivation and immunogenic context on overall tumor fitness. **C.** Illustrations demonstrating the application of the mathematical model to dissect the contributions of genetic effects, immune responses, and genotype-specific immune responses to tumor fitness using *Lkb1^KO^* as an example.

**Fig. S7:**
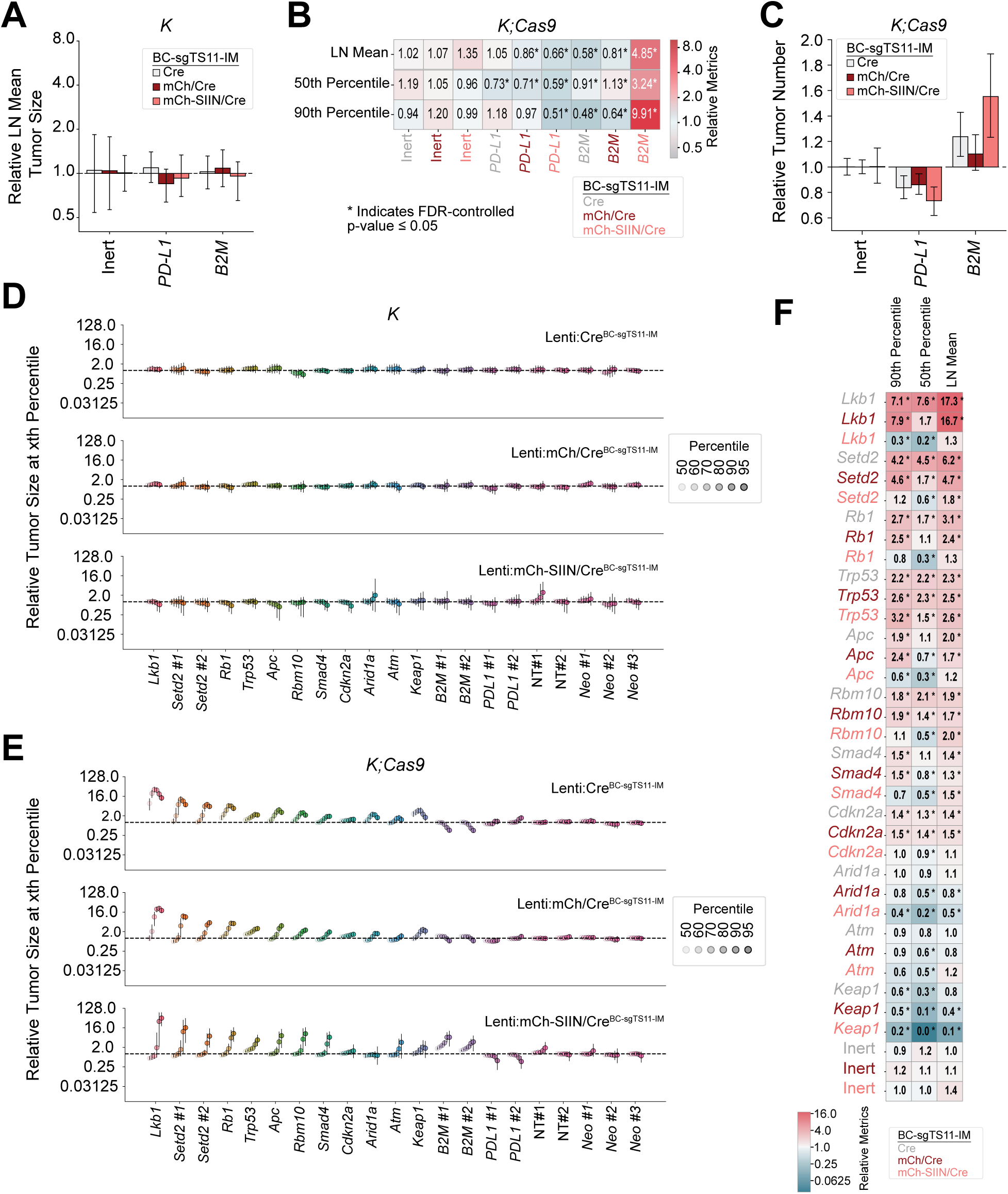
Immunogenic context shapes the effect of immunomodulatory or tumor suppressor gene inactivation on tumor outgrowth. **A.** Quantification of relative LN mean tumor size for the TS^WT,^ *PD-L1^KO^*, and *B2M^KO^* genotypes initiated with Lenti:Cre^BC-sgTS11-IM^, Lenti:mCh/Cre^BC-sgTS11-IM^, or Lenti:mCh-SIIN/Cre^BC-sgTS11-IM^ in *K* mice. Tumors were analyzed using the “Adaptive sampling method” (Fig. S6A). **B.** Heat map depicting relative tumor metrics for the TS^WT,^ *PD-L1^KO^*, and *B2M^KO^*genotypes initiated with Lenti:Cre^BC-sgTS11-IM^, Lenti:mCh/Cre^BC-sgTS11-IM^, or Lenti:mCh-SIIN/Cre^BC-sgTS11-IM^ in *K;Cas9* mice. Raw values are shown in heatmap and asterisk indicates FDR-controlled *p-*value≤ 0.05. Tumors were analyzed using the “Adaptive sampling method” (Fig. S6A). **C.** Quantification of relative tumor number for the TS^WT,^ *PD-L1^KO^*, and *B2M^KO^* genotypes initiated with Lenti:Cre^BC-sgTS11-IM^, Lenti:mCh/Cre^BC-sgTS11-IM^, or Lenti:mCh-SIIN/Cre^BC-sgTS11-IM^ in *K;Cas9* mice. Tumors were analyzed using the “Adaptive sampling method” (Fig. S6A). **D and E.** Relative tumor sizes at the indicated percentiles for each sgRNA in the BC-sgTS11-IM library in *K* (D) or *K;Cas9* (E) mice. Key indicating percentile is found in Fig. S8E (right). Tumors were analyzed using the “Adaptive sampling method” (Fig. S6A). **I. F.** Heat map depicting relative metrics in *K;Cas9* mice for tumors initiated with Lenti:Cre^BC-sgTS11-IM^, Lenti:mCh/Cre^BC-sgTS11-IM^, or Lenti:mCh-SIIN/Cre^BC-sgTS11-IM^ for all tumor suppressor genotypes. Raw values are shown in heatmap and asterisk indicates FDR-controlled *p-*value≤ 0.05. Tumors were analyzed using the “Adaptive sampling method” (Fig. S6A).

**Fig. S8:**
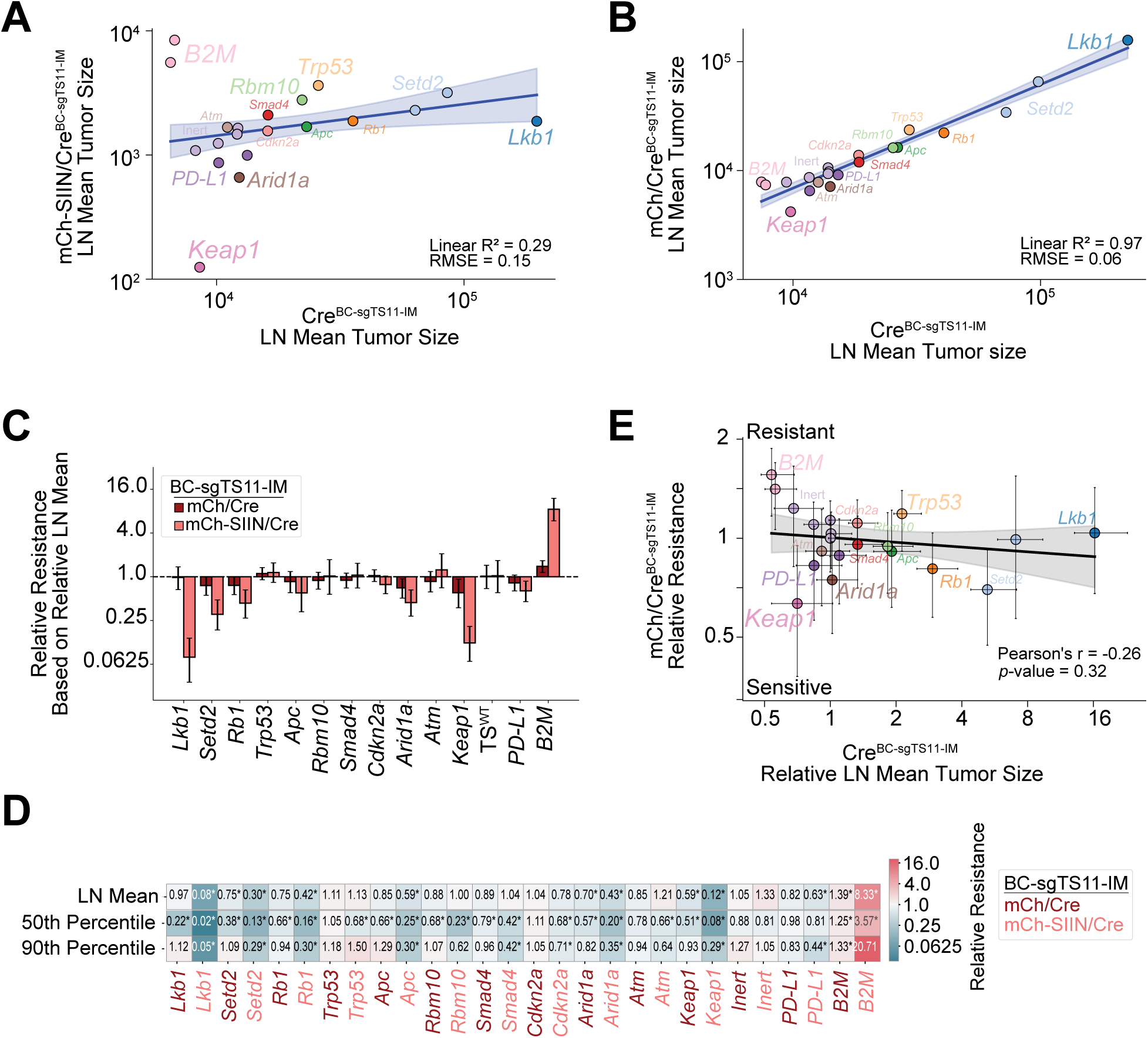
Tumor suppressor genotype modulates a tumors susceptibility to immunosurveillance upon potent neoantigen expression. **A and B.** LN mean tumor size of Cre^BC-sgTS11-IM^ tumors versus mCh-SIIN/Cre^BC-sgTS11-IM^ (A) or mCh/Cre^BC-sgTS11-IM^ (B) tumors with a linear fit. Tumors were analyzed using the “Adaptive sampling method” (Fig. S6A). **C.** Relative resistance metric calculated based on LN mean tumor size for tumors initiated by Lenti:mCh/Cre^BC-sgTS11-IM^ or Lenti:mCh-SIIN/Cre^BC-sgTS11-IM^ across each genotype. **D.** Heatmap depiction of relative resistance calculated based on different tumor metrics for tumors initiated by Lenti:mCh/Cre^BC-sgTS11-IM^ or Lenti:mCh-SIIN/Cre^BC-sgTS11-IM^ across each genotype. Raw values are shown in heatmap and asterisk indicates FDR-controlled *p-*value≤ 0.05. **E.** The relationship between the Relative Resistance metric of mCh-SIIN/Cre^BC-sgTS11-IM^ tumors and the LN mean tumor size of Lenti:Cre^BC-sgTS11-IM^ tumors for each sgRNA in the BC-sgTS11-IM pool.

**Fig. S9:**
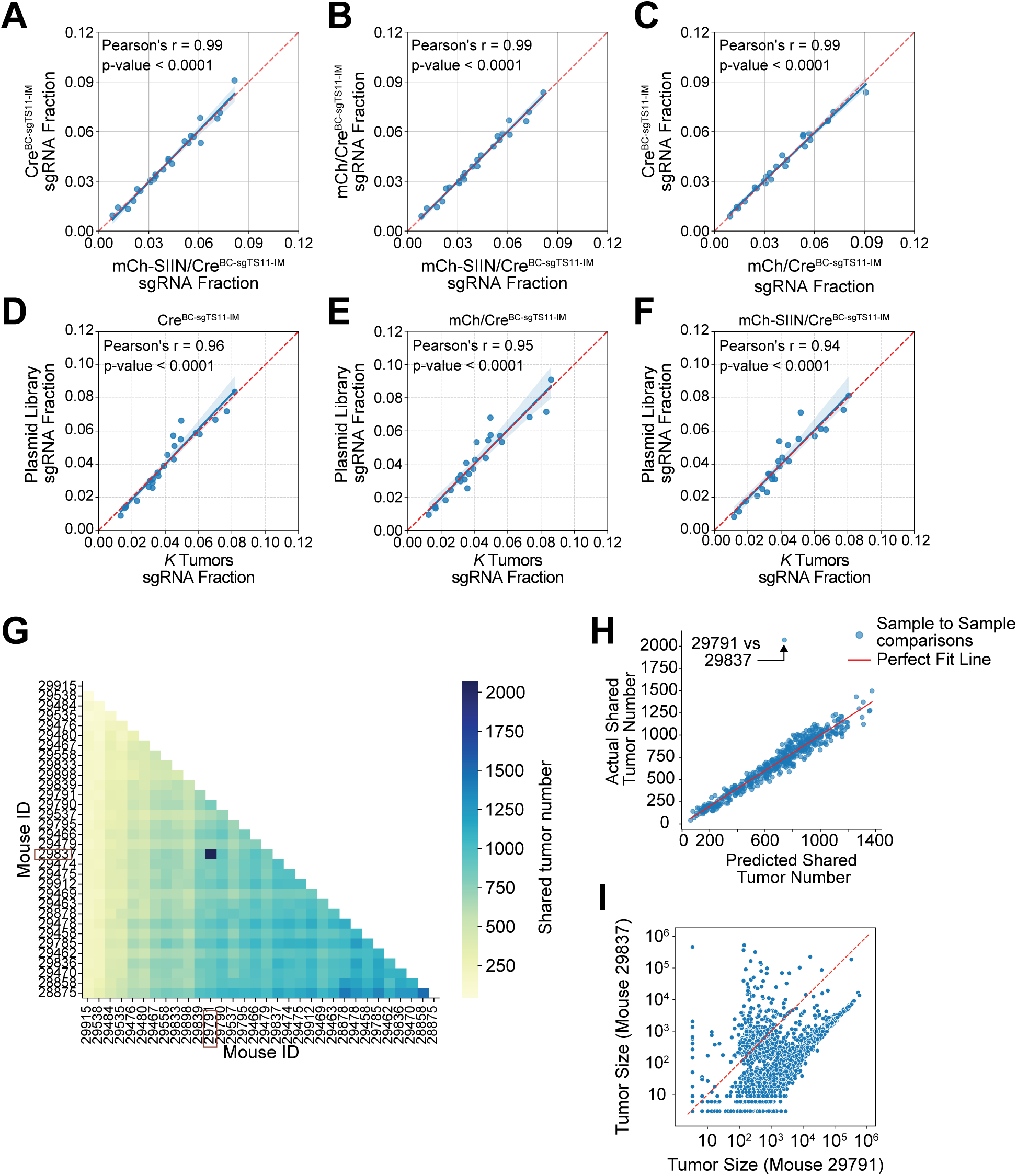
Assessing the quality of sgRNA representation and library preparation after next generation sequencing. **A to C.** Correlation of sgRNA read fractions of individual sgRNAs from each plasmid library. Each comparison is indicated in the corresponding graph. **D to F.** Correlation of sgRNA read fractions from plasmid library and *K* tumors initiated with Lenti:Cre^BC-sgTS11-IM^ (D), Lenti:mCh/Cre^BC-sgTS11-IM^ (E), or Lenti:mCh-SIIN/Cre^BC-sgTS11-IM^ (F). **G.** Heatmap of shared tumors with the same BC-sgRNA sequence in Cre^BC-sgTS11-IM^ tumors. **H.** Plot of actual versus expected shared tumor counts with the same BC-sgRNA sequence from Mouse 29791 and 29837 that identifies a higher number of shared tumors than expected due to random sampling. **I.** Tumor size from Mouse 29791 and 29837 showing a subset of tumors that aligns well with the predicted overlap value due to random sampling and another subset of tumors that indicate contamination.

